# Extraction-dependent bone proteomics reveals distinct stable and dynamic protein modules during early post-exposure degradation

**DOI:** 10.64898/2026.04.29.721604

**Authors:** Nikita Choudhary, Aparna Sajeevan, Sahla Abdulsalam, Mohammad Nasir Ahmad, Mohd. Altaf Najar

## Abstract

Bone is a highly durable biological tissue widely used in forensic, archaeological, and anthropological investigations; however, efficient protein recovery and understanding of protein stability over time remain major challenges in skeletal proteomics. Here, we systematically evaluated three bone protein extraction workflows and integrated them with data-independent acquisition (DIA) mass spectrometry to assess proteome coverage, reproducibility, and temporal protein dynamics under environmentally exposed conditions. Comparative analysis demonstrated that extraction strategy is a primary determinant of detectable proteome composition. EDTA-based demineralization followed by SDS extraction provided the deepest proteome coverage and highest reproducibility, whereas guanidine hydrochloride extraction preferentially enriched collagen and extracellular matrix proteins. In contrast, acid-based extraction yielded limited protein recovery. Temporal profiling of bone samples collected at 10 and 45 days post-exposure revealed two distinct protein classes. A temporally stable module, enriched in collagens and extracellular matrix proteins including COL1A2, COL5A2, BGN, SPARCL1, and NID2, exhibited minimal abundance change, indicating resistance to environmental degradation. In contrast, temporally dynamic proteins, enriched in mitochondrial, metabolic, and intracellular pathways such as ACO2, OGDH, PDHA1, ATP5PO, and PFKM, showed marked decline over time. These findings support a two-compartment model of bone protein preservation in which matrix-embedded proteins are preferentially retained while exposed intracellular proteins undergo progressive degradation. Collectively, this study establishes an integrated framework linking extraction methodology with temporal proteome stability and identifies candidate markers for skeletal preservation assessment and temporal biomarker development in forensic and archaeological applications.

## 1: Introduction

Bone is one of the most durable biological tissues in the vertebrate body and often remains available for investigation long after soft tissues have decomposed [1–3]. For this reason, skeletal material plays a central role in forensic identification, archaeological reconstruction, paleoanthropology, and studies of historical human remains [4, 5]. Beyond its structural importance, bone contains a complex organic matrix composed primarily of collagens together with numerous non-collagenous proteins involved in mineralization, remodeling, signaling, and cellular maintenance [1, 6]. Advances in mass spectrometry-based proteomics have increasingly enabled the characterization of this molecular landscape, creating new opportunities for biological interpretation and applied biomarker discovery [7–9].

In medicolegal investigations and research, protein-based approaches have attracted growing interest as complementary tools to DNA analysis, particularly in cases involving degraded, burned, fragmented, or environmentally exposed remains where nucleic acid recovery may be compromised [10, 11]. Bone proteins may provide information relevant to species identification, tissue origin, biological sex, age estimation, disease history, and postmortem change [12–14]. Similarly, in archaeological and paleoproteomic contexts, preserved skeletal proteins have been used to investigate taxonomy, diet, pathology, and evolutionary relationships [15–17]. These developments highlight the potential value of bone proteomics as a robust molecular resource in settings where other biomolecules are poorly preserved. Despite this promise, bone remains a challenging tissue for proteomic analysis. The mineralized hydroxyapatite matrix, extensive collagen crosslinking, and relatively low abundance of residual cellular proteins can limit extraction efficiency and reduce proteome depth [1, 18, 19]. As a result, sample preparation strategy is widely recognized as one of the most critical variables in skeletal proteomics. A variety of extraction workflows have been employed, including acid demineralization, EDTA-based decalcification, detergent-assisted lysis, and chaotropic extraction using reagents such as guanidine hydrochloride [20–22]. However, these methods often differ substantially in protein yield, selectivity, reproducibility, and compatibility with downstream mass spectrometry workflows. Direct comparative evaluation of extraction strategies for broad bone proteome recovery remains relatively limited.

An additional unresolved question concerns the temporal stability of different bone protein classes following environmental exposure. While collagen has long been recognized as a durable component of skeletal tissue, less is known about the persistence of non-collagenous extracellular matrix proteins, membrane proteins, mitochondrial proteins, metabolic enzymes, and other intracellular components [23–25]. Understanding which proteins remain stable and which decline over time is particularly relevant for forensic science, where molecular signatures of preservation or degradation may aid in estimating postmortem change, assessing sample quality, or prioritizing downstream analyses [13, 26, 27]. Moreover, distinguishing persistent proteins from degradation-sensitive proteins may improve biomarker selection for future targeted assays. Recent developments in data-independent acquisition (DIA) mass spectrometry provide an attractive platform for addressing these questions, as DIA enables high proteome coverage, improved quantitative reproducibility, and consistent detection across complex sample sets [28]. When combined with robust extraction workflows and comparative experimental design, DIA-based proteomics can provide simultaneous insight into methodological performance and biological protein stability.

In the present study, we systematically compared three bone protein extraction workflows and integrated them with DIA-based quantitative proteomics to evaluate protein recovery, reproducibility, and temporal abundance changes under environmentally exposed conditions. We further sought to identify proteins that remain stable between early and later exposure intervals, as well as proteins that show dynamic decline over time. By linking extraction methodology with temporal protein persistence, this work aims to establish a framework for future forensic, archaeological, and skeletal proteomics applications.

## 2: Materials and Methods

### 2.1: Materials

The porcine samples were collected from the Forensic Anthropology Center for Taphonomy at Yenepoya Deemed University. The cadavers were placed for the taphonomic investigations. The bones were collected from the 10-day-old and 45-day cadavers. Ethylenediaminetetraacetic acid (EDTA, cat no.E4884-100G), sodium dodecyl sulfate (SDS, Cat no. L3771-1KG), guanidine hydrochloride (GuHCl, Cat no.50940-100G), hydrochloric acid (HCl, Cat no. H1758), acetic acid (Cat no. 1.93002.2521, EMPARTA), formic acid (EMSURE, Cat no. 1.00264), Tris-HCl (Cat no. 648317-100G), and triethylammonium bicarbonate (TEAB, Cat no. T7408-500ML) used for demineralization, protein extraction (Probe Sonicator, LABMAN, L53153), and digestion were obtained from Sigma-Aldrich (St. Louis, MO, United States). Dithiothreitol (DTT, Cat no. D9779-25G), iodoacetamide (IAA, Cat no. I6125-10G), and protease inhibitor cocktail (Cat no: 4693159001) were purchased from Sigma Aldrich(St. Louis, MO, United States). Sequencing-grade modified trypsin (Cat no.V5111)was obtained from Promega (Madison, WI, United States). Pepsin (porcine gastric mucosa, Cat no. V195A) used for collagen-enriched extraction was purchased from Sigma-Aldrich (St. Louis, MO, United States). LC–MS grade water (Cat no. W6-4, 4 L), acetonitrile(Cat no.A955-4, 4L), and formic acid(Cat no. 1.00264) were obtained from Fisher Scientific (Fair Lawn, NJ, United States). Solid phase extraction disk (Cat no. 66883-U,)used in stage tip method for peptide cleanup were obtained fromThermo Fisher Scientific (Waltham, MA, United States). Protein concentration was determined using the Pierce BCA Protein Assay Kit (Cat no. 23225) and peptideconcentration was measured using the Pierce Quantitative Colorimetric Peptide Assay(Cat no. 23275) using standard spectrophotometric or colorimetric assays. Mass spectrometric analysis was performed using an Orbitrap Fusion Tribrid mass spectrometer coupled to a Vanquish Neo ultra-high-performance liquid chromatography (UHPLC) system (Thermo Fisher Scientific, Bremen, Germany). Data were acquired indata-independent acquisition (DIA) modes for protein identification and quantitative comparison across extraction workflows.

### 2.2: Methods

#### 2.2.1: Sample Collection

The bone tissue samples from the hind limb region of *Sus scrofa domesticus* were collected at two different time intervals (10 days and 45 days). The animals were euthanized using KCl (5 mE) injection intravenously with the help of the In-house Veterinary Surgeon at ASSEND Centre, Yenepoya Deemed-to-be University. After euthanasia, the animal was placed at India’s first Taphonomy Center Forensic Anthropology Center for Taphonomy (FACT) at Forensic Anthropology Unit, Department of Forensic Medicine and Toxicology, Yenepoya Medical College, Karnataka as shown in supplementary figure 1. The samples were collected from the animal’s bones with the help of veterinary surgical instruments to reach the bone site by making a knife cut through the muscles. After that, with the help of a bone rongeur (Sun Surgicals), replace it with “The bone samples were collected from the femur bones with the help of a bone rongeur (Sun Surgicals), after removing the skin and muscle tissues. A small piece, i.e., a 2 cubic cm piece of bone, was lifted out with the help of toothed forceps (5 inch). To prevent water from penetrating the bone, transparent silicone glue was used, and the muscles and skin were then sutured using autopsy twine/ligature (only at the initial stage).

The Institutional Animal Ethics Committee approved the study with a reference number of YU/IAEC/P(L)10/2024.

#### 2.2.2: Bone Sample Preparation for Proteomic Analysis

Prior to protein extraction, frozen porcine bone samples were thawed on ice and re-cleaned to remove any residual external debris. Bone fragments were rinsed with Milli-Q water, blotted dry, and transferred into pre-chilled mortar. Samples were snap-frozen in liquid nitrogen and with the help of pestle to obtain a fine homogeneous powder. Grinding under cryogenic conditions minimized heat-induced protein degradation and improved disruption of the mineralized matrix. Bone powder from each sample was thoroughly mixed, and aliquots of approximately 200 mg were transferred into 2 mL low-protein-binding microcentrifuge tubes for downstream extraction. All extraction workflows were performed in triplicate.

#### 2.2.3: Method 1: EDTA Demineralization Followed by SDS-Based Protein Extraction

For Method 1, 200 mg of bone powder was suspended in 1 mL of 0.5 M EDTA (pH 8.0) supplemented with protease inhibitor cocktail. Samples were incubated at 4°C on an end-over-end rotator for 48 h to achieve demineralization. Fresh EDTA solution was replaced after 24 h to improve calcium chelation efficiency. Following incubation, samples were centrifuged at 15,000 × g for 10 min at 4°C, and the supernatant was discarded. The demineralized pellet was washed twice with cold Milli-Q water to remove residual EDTA.Protein extraction was then performed by adding 500 µL of lysis buffer containing 4% SDS, 100 mM Tris-HCl (pH 8.0), and 10 mM DTT. Samples were vortexed thoroughly and heated at 95°C for 10 min to denature proteins and disrupt collagen-associated crosslinks. Tubes were then incubated for 30 min at room temperature with intermittent agitation. Where required, brief sonication was applied to improve solubilization. Insoluble debris was removed by centrifugation at 16,000 × g for 15 min, and the clarified supernatant containing extracted proteins was collected for downstream processing.

#### 2.2.4: Method 2: EDTA Demineralization Followed by Guanidine Hydrochloride Extraction

For Method 2, 200 mg of bone powder was mixed with 1 mL of 0.5 M EDTA (pH 8.0) containing protease inhibitors and incubated at 4°C for 24 h with continuous rotation. Compared with Method 1, a shorter demineralization step was used to reduce loss of loosely associated extracellular matrix proteins. Samples were centrifuged at 15,000 × g for 10 min, and the supernatant was removed. Pellets were washed twice with cold Milli-Q water.The demineralized pellet was then extracted in 1 mL of chaotropic buffer containing 4 M guanidine hydrochloride and 50 mM Tris-HCl (pH 7.4) supplemented with protease inhibitors. Samples were vortexed vigorously and incubated overnight (12–16 h) at 4°C on a rotator. Mild sonication was applied when necessary to enhance extraction efficiency. Following incubation, samples were centrifuged at 16,000 × g for 20 min at 4°C, and the supernatant containing solubilized proteins was collected. Guanidine hydrochloride was subsequently removed during downstream cleanup prior to digestion.

#### 2.2.5: Method 3: Acid Demineralization Followed by Pepsin-Assisted Extraction

For Method 3, 200 mg of bone powder was suspended in 1 mL of 0.6 M HCl and incubated at 4°C for 24 h with gentle rotation to dissolve the mineral phase. Following centrifugation at 15,000 × g for 10 min, the acid supernatant was discarded and the pellet was washed two to three times with cold Milli-Q water until near-neutral pH was achieved.The washed pellet was resuspended in 1 mL of extraction buffer consisting of 0.5 M acetic acid containing pepsin (1 mg/mL). Samples were incubated overnight (16–24 h) at 4°C with rotation to facilitate collagen-enriched solubilization. Following extraction, tubes were centrifuged at 16,000 × g for 20 min and the supernatant was collected. Extracts were slowly neutralized using 1 M Tris base prior to reduction, alkylation, and enzymatic digestion.

#### 2.2.5: Protein Reduction, Alkylation, and Enzymatic Digestion

For all extraction workflows, protein extracts were reduced with DTT to a final concentration of 10 mM at 56°C for 30 min. Samples were then cooled to room temperature and alkylated with iodoacetamide to a final concentration of 20 mM for 30 min in the dark. Excess iodoacetamide was quenched by addition of DTT.Samples containing SDS were processed using S-Trap microcolumns or filter-aided sample preparation (FASP) according to manufacturer recommendations. Guanidine-containing extracts were subjected to acetone precipitation prior to digestion. Protein pellets were resuspended in 100 mM TEAB buffer. Sequencing-grade trypsin was added at an enzyme-to-protein ratio of 1:50, and digestion was performed overnight at 37°C. Reactions were terminated by acidification with formic acid.

#### 2.2.6: Peptide Cleanup and LC–MS Preparation

Tryptic peptides were desalted using C18 solid-phase extraction cartridges or StageTips, washed to remove salts and contaminants, and eluted using 50% acetonitrile containing 0.1% formic acid. Eluted peptides were dried under vacuum in a SpeedVac concentrator and reconstituted in 0.1% formic acid prior to LC–MS/MS analysis. Peptide concentration was determined using spectrophotometric or colorimetric assays where required.

#### 2.2.7: LC–MS/MS Data Acquisition

Peptide samples were analyzed using an Orbitrap Fusion Tribrid mass spectrometer coupled online to a Vanquish Neo ultra-high-performance liquid chromatography (UHPLC) system (Thermo Fisher Scientific, Bremen, Germany). Peptides were separated by reversed-phase nanoLC and introduced into the mass spectrometer through a nano-electrospray ionization source operated in positive ion mode. Mobile phase A consisted of water containing 0.1% formic acid, and mobile phase B consisted of 80% acetonitrile containing 0.1% formic acid. Peptides were loaded onto the analytical column and separated at a constant flow rate of 300 nL/min using a 120 min gradient. The chromatographic program was as follows: initial equilibration at 5% B, followed by a linear increase to 40% B over 90 min, ramping to 100% B over 15 min, held at 100% B for 5 min for column washing, followed by re-equilibration at 2–5% B prior to the next injection. Total run time for each sample was 120 min. The nano-electrospray ionization source was operated using a spray voltage of 2.1 kV. Ion transfer tube temperature was maintained at 275°C. The instrument was operated in liquid chromatography mode with automatic peak detection enabled. Internal mass calibration was disabled during acquisition. Survey full-scan mass spectra (MS1) were acquired in the Orbitrap analyzer at a resolution of 120,000 (at m/z 200). The scan range was set from m/z 350–1600. RF lens was operated at 55%, and the maximum injection time was 20 ms. Precursor ions with charge states of 2+ or higher were considered for fragmentation. Tandem mass spectra (MS/MS) were acquired using higher-energy collisional dissociation (HCD) with Orbitrap detection. Precursor ions were isolated using an isolation window of 20 m/z. Fragmentation was performed using a normalized collision energy of 35%. MS/MS spectra were acquired in the Orbitrap at a resolution of 15,000 over a scan range of m/z 145–2000. RF lens was set to 40%. The instrument was operated using a time-controlled duty cycle of 5 s to maximize peptide sampling across chromatographic peaks. For comparative quantitative proteomics, the instrument was operated in data-independent acquisition (DIA) mode, enabling reproducible fragmentation of precursor windows across the chromatographic run. This approach provided consistent peptide quantification across multiple biological replicates, extraction methods, and temporal exposure groups.

#### 2.2.8: Proteomic Data Processing and Database Searching Using DIA-NN

Raw mass spectrometry files generated from DIA acquisitions were processed using DIA-NN (Data-Independent Acquisition by Neural Networks; version 2.3.1or latest available release) for peptide identification, spectral matching, and label-free quantitative analysis. All files were searched against the Sus scrofa (pig) reference proteome downloaded from the UniProt Knowledgebase (UniProtKB), containing reviewed and annotated protein entries available at the time of analysis. A common contaminants database including keratins, trypsin, and laboratory background proteins was appended to the FASTA file, and decoy sequences were generated automatically by DIA-NN for false discovery rate estimation. Searches were performed assuming tryptic specificity, with cleavage after lysine and arginine residues except when followed by proline, allowing up to two missed cleavages. Carbamidomethylation of cysteine was specified as a fixed modification, while oxidation of methionine, acetylation of protein N-termini, and acetylation of lysine residues were included as variable modifications. Precursor and fragment mass tolerances were automatically optimized by DIA-NN according to instrument mass accuracy. Retention time alignment across runs, neural network-assisted peak scoring, and interference correction were enabled to improve peptide detection and quantitative consistency between samples. Library-basedsearching was performed using DIA-NN predicted spectra and retention time models generated directly from the FASTA database, with iterative refinement of in silico libraries where applicable. Peptide precursors, peptides, and protein groups were filtered at a 1% false discovery rate (FDR) using the target-decoy strategy. Protein quantification was based on aggregated precursor ion intensities, and cross-run normalization was enabled to minimize technical variation. In addition, bovine serum albumin (BSA) spiked during sample preparation was used as an internal reference for normalization where required. Final precursor-, peptide-, and protein-level quantitative output tables were exported from DIA-NN and further analyzed using R statistical software for log2 transformation, missing-value filtering, principal component analysis, hierarchical clustering, differential abundance testing, volcano plot generation, temporal stability analysis, and functional enrichment analysis.

#### 2.2.9: Bioinformatics and Statistical Analysis

Protein group intensity data generated from DIA-NN were used for downstream bioinformatics analysis and visualization. Raw intensity values were log2-transformed to stabilize variance and improve comparability across samples. To reduce technical variability between runs, cross-run normalization performed within DIA-NN was supplemented by internal normalization using bovine serum albumin (BSA) spiked during sample preparation. Proteins with inconsistent detection or excessive missing values across replicates were filtered to retain high-confidence quantitative data for further analysis. Exploratory data analysis and visualization were carried out using a combination of web-based and statistical tools. Heatmaps, hierarchical clustering, and k-means clustering were generated using Morpheus (https://software.broadinstitute.org/morpheus/) to identify patterns of protein abundance across extraction methods and time points. Venn diagram analysis to assess protein overlap between datasets was performed using BioVenn (https://www.biovenn.nl/index.php). Functional enrichment analysis was conducted using ShinyGO (version 0.85.1) (https://bioinformatics.sdstate.edu/go/) to identify significantly enriched Gene Ontology (GO) categories, including biological processes, molecular functions, and cellular components. Enrichment results were filtered based on statistical significance thresholds, and relevant pathways were visualized accordingly. Additional statistical analyses, including data filtering, transformation, correlation analysis, principal component analysis (PCA), and graphical visualization (e.g., violin plots and clustering outputs), were performed using R Studio (R Foundation for Statistical Computing). Based on fold change and statistical criteria, proteins were categorized into temporally stable and temporally dynamic groups for downstream biological interpretation.

## 3: Results

To systematically characterize the bone proteome and evaluate the impact of extraction strategies, we performed a comprehensive proteomic analysis using multiple extraction methods followed by quantitative mass spectrometry. Comparative analysis revealed significant differences in protein recovery across methods, with distinct subsets of proteins identified under each condition. Subsequent analyses focused on datasets generated using the most effective extraction approaches, enabling robust assessment of data quality, temporal variation, and functional organization of the bone proteome. Notably, our results revealed the presence of both temporally stable and dynamically changing protein populations, reflecting distinct biological and structural properties within bone tissue.

### 3.1: Comparative evaluation of extraction strategies demonstrates distinct protein recovery profiles

A comprehensive workflow for bone protein extraction and DIA-based proteomic analysis is illustrated in Figure 1. To systematically assess the impact of extraction methodology on proteome coverage, three distinct approaches EDTA-based demineralization with SDS lysis (Method 1), guanidine hydrochloride-based extraction (Method 2), and acid demineralization coupled with pepsin digestion (Method 3) were evaluated across biological replicates. Quantitative comparison of protein identifications revealed substantial differences in extraction efficiency among the three methods. Across replicates, Method 1 consistently yielded the highest number of identified proteins, with 1321, 1269, and 1242 proteins detected in replicates R1, R2, and R3, respectively (Table 1). In contrast, Method 2 demonstrated moderate proteome coverage, with protein identifications ranging from 854 to 1088 across replicates. Method 3 showed markedly reduced protein recovery, with fewer than 150 proteins identified per replicate, indicating limited suitability for comprehensive bone proteome profiling. Analysis of reproducibility across replicates further highlighted the robustness of Method 1, which exhibited 989 proteins consistently identified across all three replicates, compared to 728 proteins for Method 2 and only 67 proteins for Method 3 (Table 1). These results indicate that Method 1 not only provides broader proteome coverage but also demonstrates superior reproducibility.

**Figure 1.**
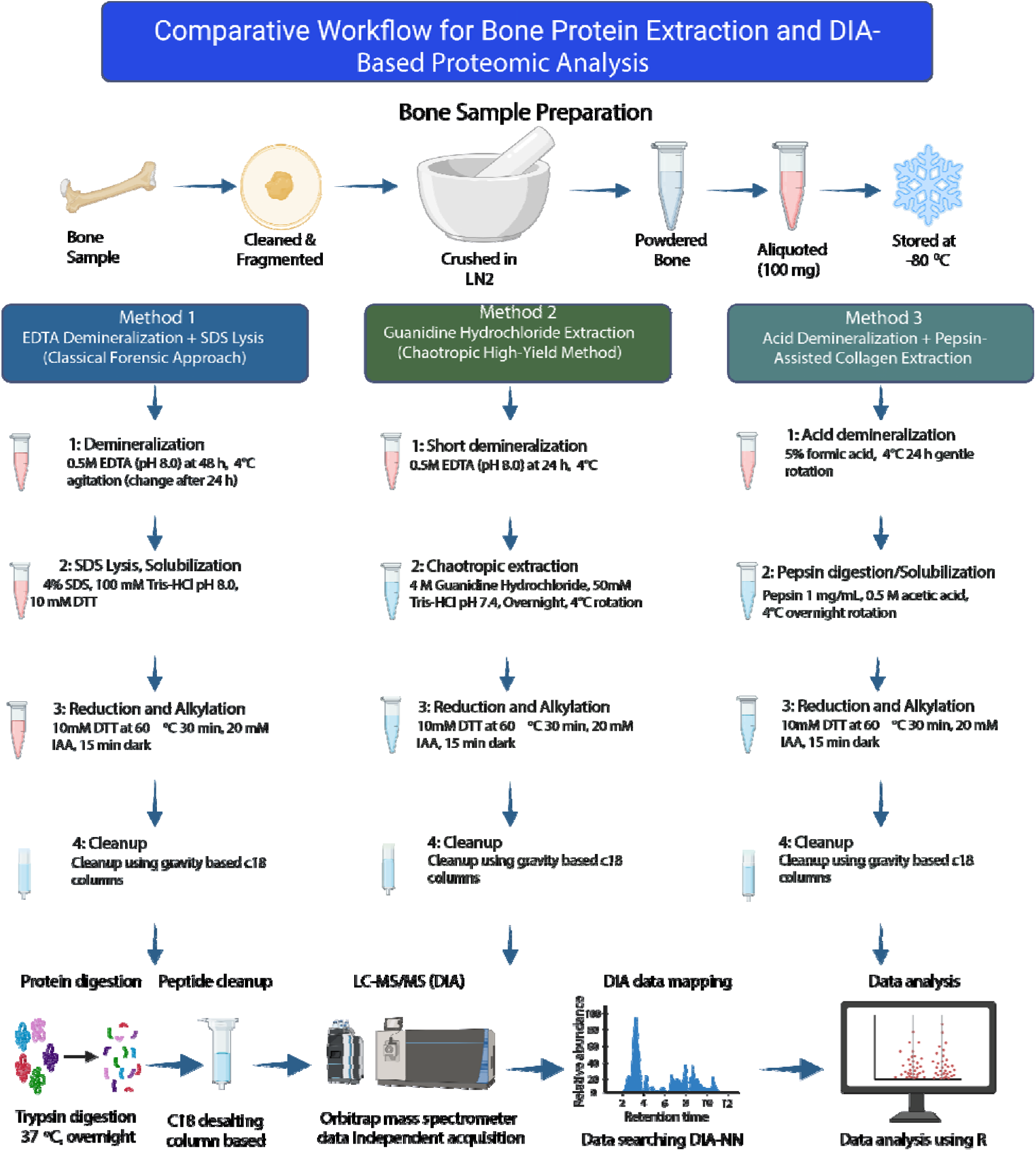
Comparative workflow for bone protein extraction and DIA-based proteomic analysis: Schematic representation of the experimental workflow used for bone proteomics. Bone samples were cleaned, fragmented, cryogenically powdered, and aliquoted prior to protein extraction. Three extraction strategies were evaluated: (i) EDTA-based demineralization followed by SDS lysis (Method 1), (ii) guanidine hydrochloride-based chaotropic extraction (Method 2), and (iii) acid demineralization coupled with pepsin digestion (Method 3). Extracted proteins were subjected to reduction, alkylation, and C18-based cleanup, followed by tryptic digestion. Peptides were analyzed using LC–MS/MS in data-independent acquisition (DIA) mode, and protein identification and quantification were performed using DIA-NN, followed by downstream data analysis.

**Table 1:**
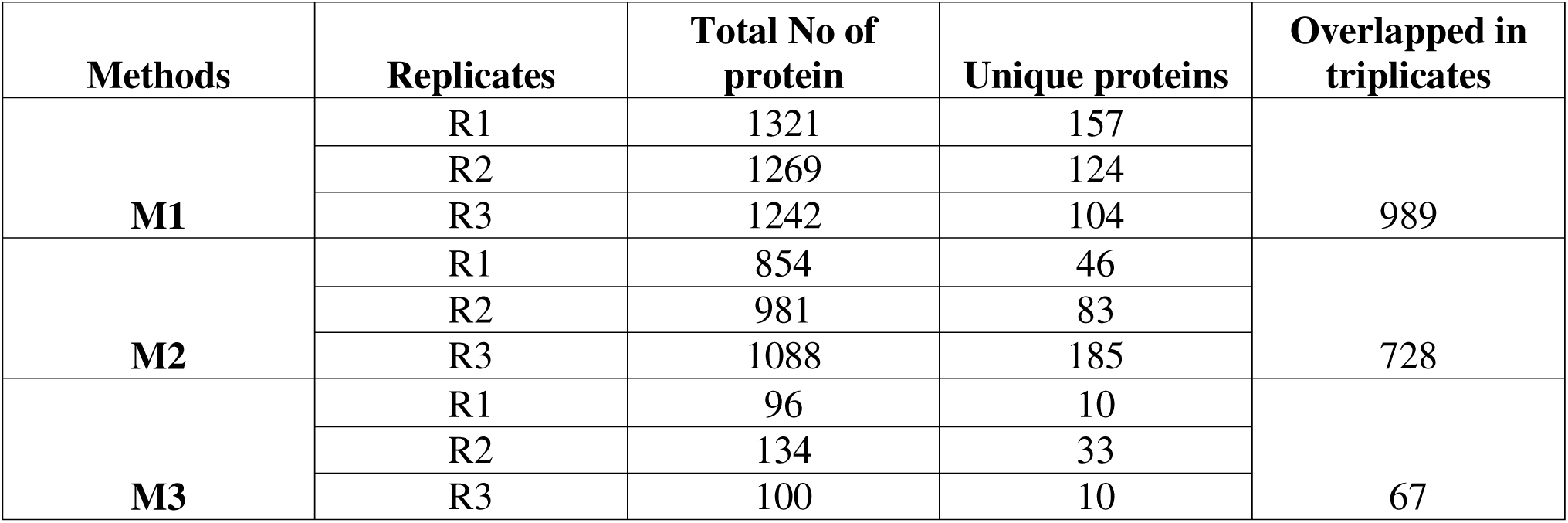
Summary of protein identifications and reproducibility across extraction methods.

### 3.2: Overlap analysis highlights method-dependent proteome capture

To further investigate similarities and differences in protein recovery across replicates, overlap analysis was performed using Venn diagrams (Figure 2, Supplementary Figure 2). Method 1 showed a substantial core proteome shared across replicates, with a large number of overlapping proteins, alongside a moderate number of uniquely identified proteins in individual replicates. Method 2 exhibited a similar but reduced overlap pattern, indicating partial consistency in protein identification. In contrast, Method 3 displayed minimal overlap across replicates, reflecting both low protein yield and limited reproducibility. Importantly, Method 1 demonstrated the highest number of unique protein identifications across replicates, suggesting enhanced sensitivity in capturing a broader range of proteins. Method 2 contributed additional unique proteins not detected by Method 1, indicating complementary proteome coverage between these two methods. However, Method 3 contributed minimally to overall protein identification, further supporting its limited applicability for in-depth proteomic analysis.

**Figure 2.**
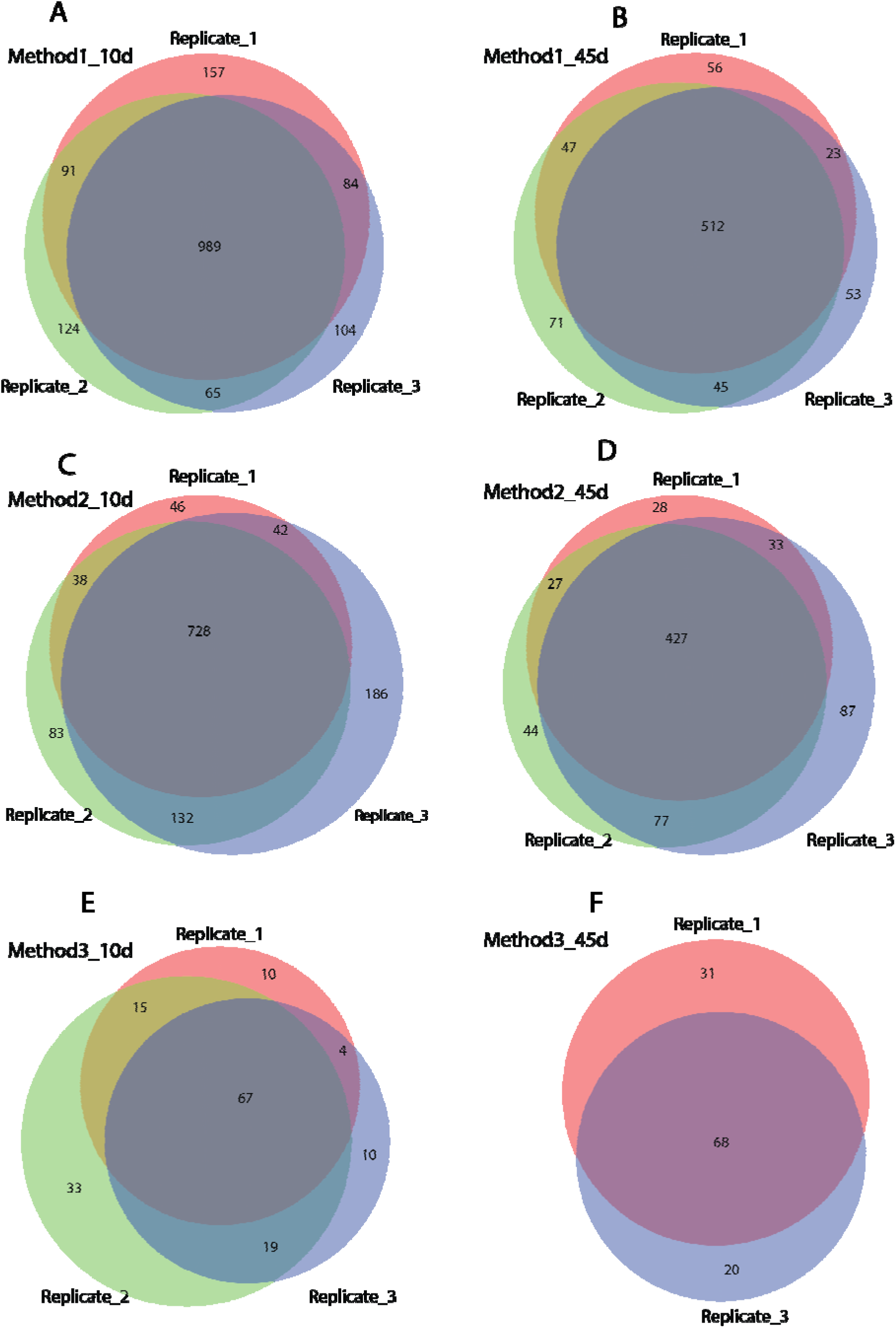
Comparison of protein identification across extraction methods and replicates: Venn diagrams illustrating the overlap and unique protein identifications across biological replicates for each extraction method. (A) Method 1 (EDTA + SDS) shows a high degree of overlap across replicates with a large shared core proteome. (B) Method 2 (guanidine hydrochloride) demonstrates moderate overlap with both shared and unique protein identifications. (C) Method 3 (acid demineralization + pepsin) exhibits limited protein identification and reduced overlap across replicates. Numbers within each region indicate the number of proteins identified uniquely or shared among replicates.

### 3.3: Temporal comparison reveals method-dependent protein retention

To assess the consistency of protein detection across time points, protein identifications at 10-day and 45-day intervals were compared for Methods 1 and 2 (Table 2). Method 1 identified a total of 1614 proteins at 10 days and 807 proteins at 45 days, with 662 proteins overlapping between the two time points. Similarly, Method 2 identified 1255 proteins at 10 days and 723 proteins at 45 days, with 570 proteins shared across time points.

**Table 2:**
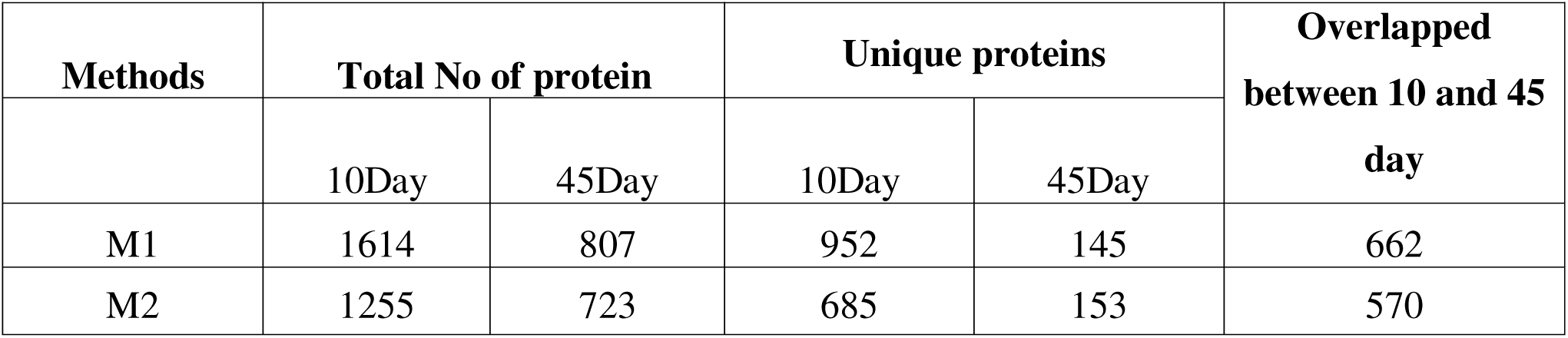
Comparison of protein identifications across time points for Methods 1 and 2.

| Methods | Total No of protein |  | Unique proteins |  | Overlapped<br>between 10 and 45<br>day |
| --- | --- | --- | --- | --- | --- |
|  | 10Day | 45Day | 10Day | 45Day |  |
| M1 | 1614 | 807 | 952 | 145 | 662 |
| M2 | 1255 | 723 | 685 | 153 | 570 |

Notably, both methods exhibited a reduction in total protein identifications at the later time point, suggesting time-dependent loss of detectable proteins. However, a substantial subset of proteins remained consistently detected across both time points, indicating the presence of a stable core proteome. Method 1 exhibited a larger overlap between time points compared to Method 2, further supporting its superior performance in maintaining proteome coverage over time. Taken together, these results demonstrate that Method 1 provides the most comprehensive and reproducible proteome coverage, while Method 2 offers complementary protein identification with moderate efficiency. In contrast, Method 3 exhibits limited protein recovery and poor reproducibility, rendering it unsuitable for detailed proteomic characterization of bone samples. Based on these findings, all subsequent analyses were performed using datasets generated from Method 1 and Method 2.

### 3.4: Differential recovery of collagen proteins across extraction methods

Given that collagens represent the dominant structural components of bone extracellular matrix [29], we next assessed the efficiency of each extraction method in recovering collagen proteins. Quantitative analysis revealed notable differences in both the number of collagen proteins identified and the depth of peptide coverage across methods (Table 3, Supplementary Table S1).

**Table 3:** Summary of collagen protein identification across extraction methods.

| Methods | Total No of protein | Peptides |
| --- | --- | --- |
| M1 | 73 | 1625 |
| M2 | 79 | 1817 |
| M3 | 29 | 162 |

Method 2 resulted in the highest number of identified collagen proteins (79) and exhibited the greatest peptide coverage (1817 peptides), indicating enhanced extraction efficiency for extracellular matrix components. Method 1 identified a comparable number of collagen proteins (73 proteins) with substantial peptide coverage (1625 peptides), demonstrating robust performance in recovering structurally embedded matrix proteins. In contrast, Method 3 showed markedly reduced collagen identification, with only 29 collagen proteins and significantly lower peptide coverage (162 peptides), reflecting limited extraction efficiency.

Detailed analysis of collagen subtypes further highlighted differences in method performance. Both Method 1 and Method 2 enabled the identification of major fibrillar collagens, including type I (COL1A1, COL1A2), type III (COL3A1), and type II (COL2A1), as well as multiple non-fibrillar collagens such as type IV (COL4A1, COL4A2) and type VI (COL6A1, COL6A2, COL6A3) (Supplementary Table S1). Notably, Method 2 demonstrated broader coverage of less abundant collagen types, including COL18A1, COL22A1, and COL24A1, suggesting improved sensitivity toward diverse extracellular matrix components.

In addition to structural collagens, several collagen-associated proteins involved in matrix remodeling and processing, including MMP2, MMP13, and PCOLCE, were also identified, further supporting effective extraction of extracellular matrix-associated proteins.

Overall, these findings indicate that while Method 1 provides comprehensive proteome coverage, Method 2 demonstrates enhanced specificity and depth for collagen and extracellular matrix protein recovery, whereas Method 3 exhibits limited capability in extracting bone matrix components. This highlights the importance of extraction strategy in capturing biologically relevant extracellular matrix proteins in bone proteomics.

### 3.4: Normalization and multivariate analyses demonstrate high data quality, strong reproducibility, and biologically meaningful variation

To enable reliable quantitative comparison across all samples, protein abundance values were normalized using bovine serum albumin (BSA) as an internal spike-in standard introduced during sample preparation. This strategy was employed to minimize variability arising from sample handling, digestion efficiency, and instrument response, thereby improving comparability across extraction methods, biological replicates, and temporal conditions. Following normalization, samples were analyzed collectively to assess overall dataset quality and the principal sources of variation. Global visualization of normalized protein abundance profiles revealed clear and structured expression patterns across the dataset. Unsupervised heatmap analysis with k-means clustering identified multiple groups of proteins displaying coordinated abundance trends across samples (Figure 3A). Several clusters exhibited method-associated enrichment patterns, whereas others showed temporal abundance shifts between 10-day and 45-day samples. These findings indicate that the dataset captures systematic biological differences rather than random technical noise.

**Figure 3.**
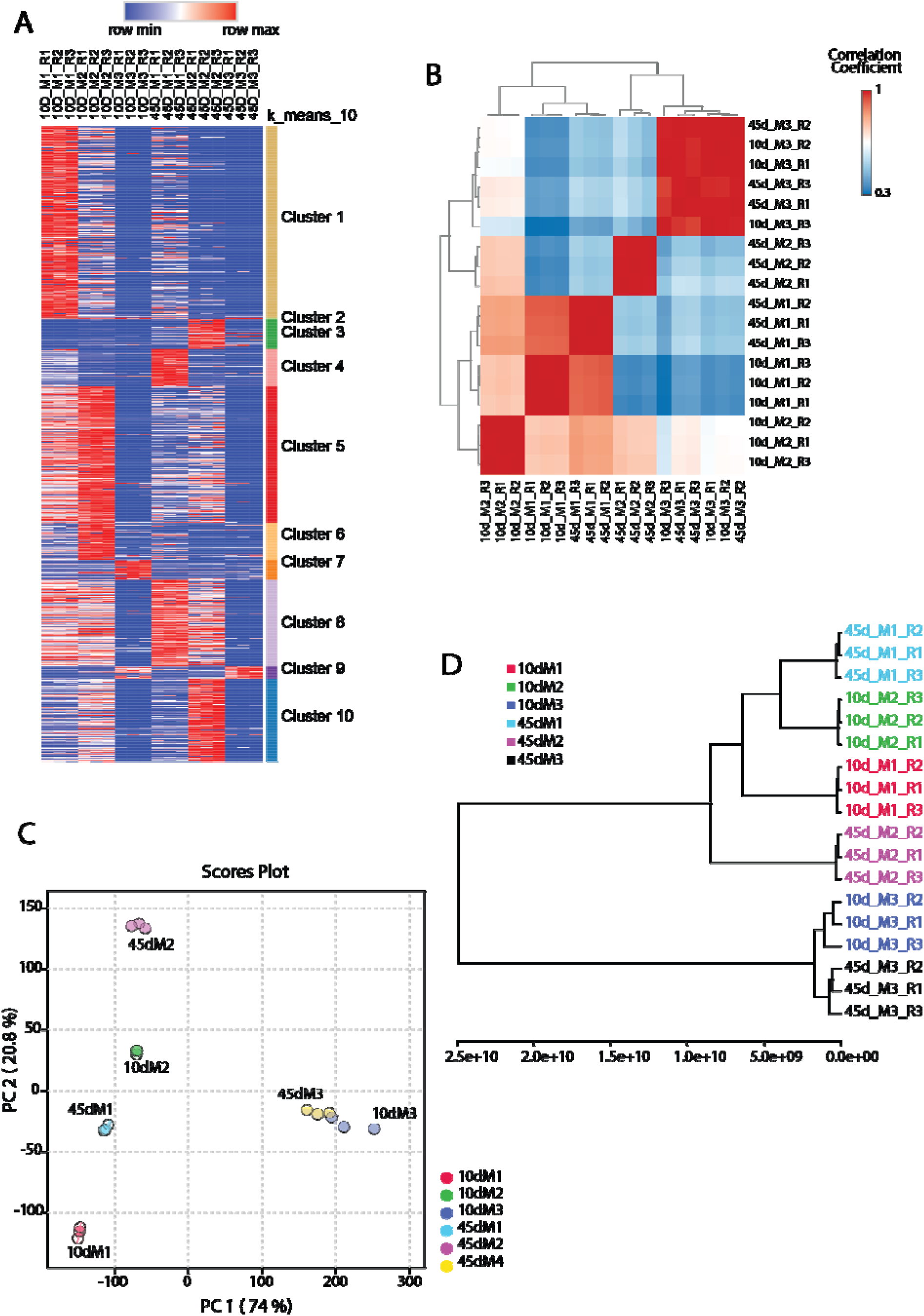
Global assessment of quantitative data quality, reproducibility, and sample relationships: **(A)** Heatmap of normalized protein abundance values across all samples with k-means clustering, showing coordinated protein expression patterns and distinct abundance modules across methods and time points. **(B)** Pairwise correlation matrix based on normalized protein intensities demonstrating strong reproducibility among biological replicates and separation between experimental conditions.**(C)** Principal component analysis (PCA) of normalized quantitative proteomic data showing clustering of replicates and separation of samples according to extraction method and temporal condition (10-day vs 45-day).**(D)** Hierarchical clustering dendrogram of samples based on global protein abundance profiles, illustrating grouping primarily by extraction method and secondarily by time point.

To further assess quantitative consistency, pairwise sample correlation analysis was performed using normalized protein intensities (Figure 3B). Biological replicates from the same experimental condition demonstrated high correlation coefficients, reflecting strong analytical reproducibility and stable quantification across independent sample preparations. In contrast, lower correlation values were observed between samples processed using different extraction methods or collected at different time points, indicating that method-dependent recovery and temporal changes represent major contributors to global proteomic variation.

Principal component analysis (PCA) was next used to define the dominant axes of variance within the dataset (Figure 3C). The first principal component (PC1), explaining approximately 74% of the total variance, clearly separated samples according to extraction strategy, demonstrating that protein recovery methodology exerts a major influence on the detectable bone proteome. Samples extracted using Method 1 and Method 2 occupied distinct regions of the PCA space, consistent with differences in protein composition and extraction selectivity. The second principal component (PC2), accounting for approximately 20.8% of the variance, further contributed to separation between 10-day and 45-day samples within extraction groups. This pattern suggests that temporal exposure influences the abundance of a subset of proteins independent of extraction strategy, consistent with progressive remodeling or degradation of specific bone-associated proteins over time.

Importantly, replicate samples clustered tightly within each condition in PCA space, indicating minimal technical dispersion relative to the observed biological differences. The close grouping of replicates provides strong evidence for reproducible sample processing, stable LC–MS/MS performance, and reliable quantitative normalization. Hierarchical clustering of samples based on global protein abundance profiles independently confirmed these observations (Figure 3D). Samples grouped primarily according to extraction method, followed by secondary clustering according to temporal condition, while replicates consistently clustered together. This concordance between PCA, correlation analysis, and hierarchical clustering further supports the robustness of the dataset. In addition to multivariate analyses, direct pairwise comparison between 45-day and 10-day samples using differential abundance analysis further supported the presence of temporal proteomic remodeling. Volcano plot analysis identified multiple significantly altered proteins in both Method 1 and Method 2 datasets, with proteins showing both increased and decreased abundance at 45 days relative to 10 days (Supplementary Figure 3). Notably, both methods exhibited a subset of strongly changing proteins, indicating that temporal exposure influences specific components of the bone proteome while a large proportion of proteins remain comparatively stable.

Collectively, these results demonstrate that the quantitative proteomic workflow generated a high-quality and internally consistent dataset in which the dominant sources of variation are attributable to extraction methodology and temporal differences rather than technical noise. These findings validate the suitability of the dataset for downstream comparative analyses aimed at identifying temporally stable and dynamically changing protein populations in bone.

### 3.5: Identification of temporally stable protein modules reveals persistent bone matrix components

To identify proteins that remain consistently detectable following prolonged environmental exposure, normalized abundance profiles were analyzed using unsupervised clustering across all samples. Heatmap analysis of the combined quantitative dataset identified ten major abundance clusters with distinct expression patterns across extraction methods and time points (Figure 4A). Among these clusters, Cluster 8 was of particular interest, as proteins within this group displayed highly similar abundance profiles across both 10-day and 45-day samples and across Method 1 and Method 2 extraction workflows. In contrast to dynamically changing clusters, proteins within Cluster 8 exhibited minimal temporal variation, suggesting relative resistance to degradation and stable retention within bone tissue. To further validate this pattern, proteins belonging to Cluster 8 were re-clustered separately (Figure 4B). The expanded heatmap confirmed a conserved abundance pattern across biological replicates and experimental groups, supporting the existence of a robust temporally stable protein module. Quantitative distribution of Cluster 8 protein abundances was further assessed by violin plot analysis (Figure 4C), which demonstrated highly comparable abundance distributions across all sample groups, with no substantial shift between 10-day and 45-day conditions. These findings indicate that proteins within this cluster are minimally affected by temporal exposure or extraction strategy.

**Figure 4.**
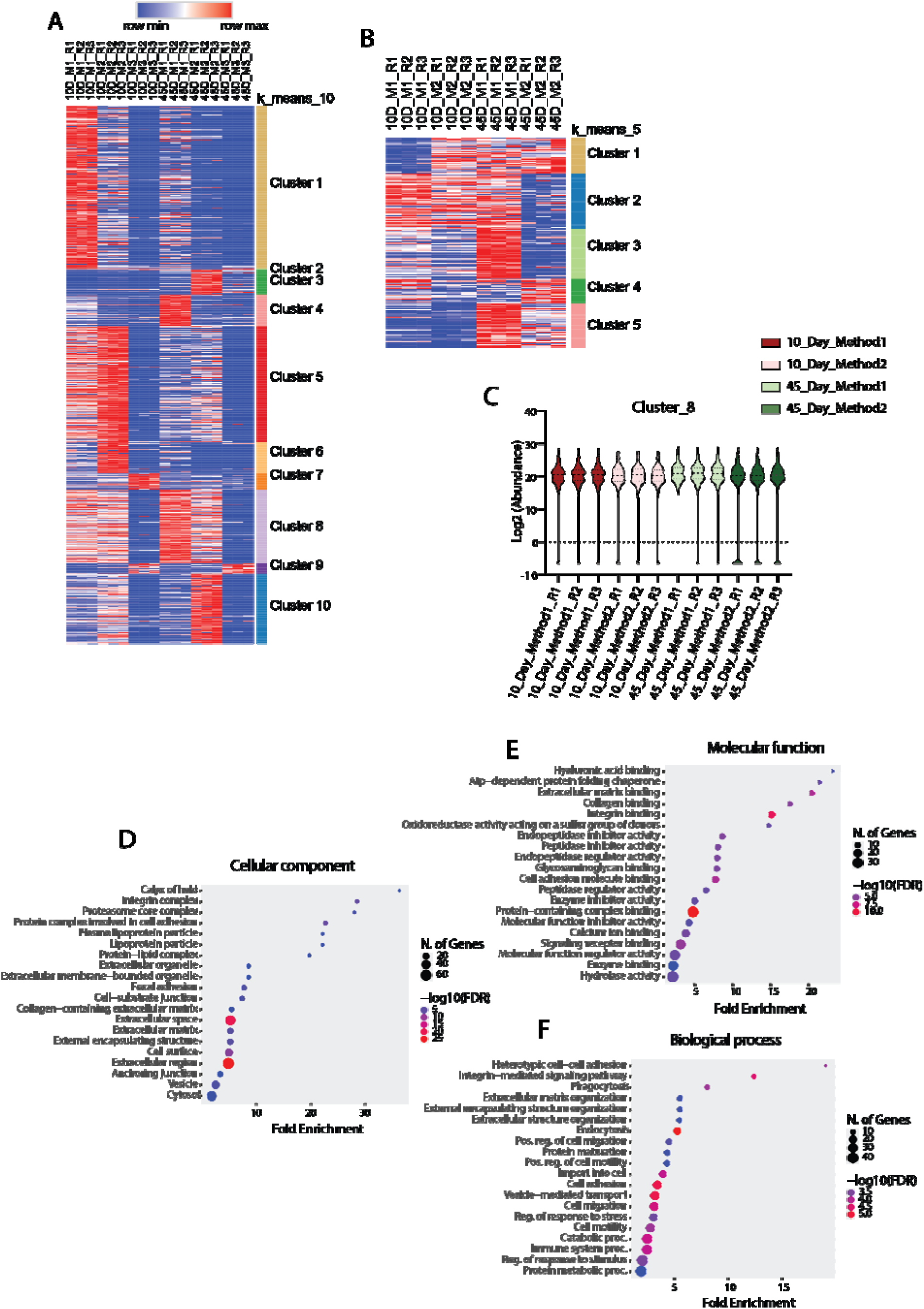
Identification and functional characterization of a temporally stable bone protein module: **(A)** Heatmap of normalized abundance values for all quantified proteins across samples, showing ten major abundance clusters identified by k-means clustering.**(B)** Expanded heatmap of Cluster 8 proteins demonstrating highly consistent abundance across extraction methods and time points.**(C)** Violin plot showing log2-transformed abundance distribution of Cluster 8 proteins across all sample groups, indicating minimal temporal variation.**(D)** Gene Ontology cellular component enrichment analysis of Cluster 8 proteins.**(E)** Gene Ontology molecular function enrichment analysis of Cluster 8 proteins.**(F)** Gene Ontology biological process enrichment analysis of Cluster 8 proteins

#### 3.5.1: High-confidence stable proteins remain unchanged across time

To identify the most persistent candidates within this stable module, a more stringent filtering strategy was applied to select proteins showing minimal fold change and non-significant temporal differences across conditions. This analysis identified a subset of high-confidence temporally stable proteins, hereafter referred to as super-stable proteins (Supplementary Figure 4A–D). These proteins exhibited highly conserved abundance patterns across replicates, methods, and time points, further supporting their resistance to degradation. The presence of statistically unchanged proteins across prolonged exposure conditions highlights their potential utility as robust reference markers for quantitative normalization, skeletal tissue characterization, and forensic biomarker development.

#### 3.5.2: Stable proteins are enriched in extracellular matrix and structural pathways

To determine the biological identity of the stable protein module, Gene Ontology enrichment analysis was performed on Cluster 8 proteins. Cellular component analysis revealed strong enrichment for extracellular matrix, collagen-containing extracellular matrix, extracellular region, anchoring junction, focal adhesion, and cell surface terms (Figure 4D), indicating preferential localization to structurally protected extracellular compartments. Molecular function analysis identified enrichment of collagen binding, integrin binding, calcium ion binding, protein-containing complex binding, and enzyme inhibitor activity (Figure 4E), consistent with roles in matrix organization and extracellular structural integrity. Biological process enrichment demonstrated association with extracellular matrix organization, cell adhesion, regulation of motility, protein maturation, and response to stimulus (Figure 4F), supporting a functional role in tissue architecture and matrix maintenance. Collectively, these results indicate that temporally stable proteins are predominantly extracellular and matrix-associated, suggesting that physical embedding within the mineralized bone matrix contributes to their long-term persistence.

The persistence of these proteins across time points suggests reduced susceptibility to environmental degradation relative to intracellular or metabolically active proteins. Their structural association with the extracellular matrix likely provides protection from proteolysis and chemical degradation. Accordingly, these proteins represent promising candidates for forensic persistence markers in aged or environmentally exposed skeletal remains.

### 3.6: Temporally stable proteins identify candidate forensic persistence markers

To refine the broader stable protein module into markers of potential forensic relevance, a subset of temporally stable proteins was selected from Method 1 based on three criteria: (i) minimal abundance change between 10-day and 45-day samples, (ii) known association with bone extracellular matrix or skeletal biology, and (iii) robust quantitative detection across replicates. Quantitative comparison of these proteins confirmed highly consistent abundance profiles across both time points, supporting their resistance to early environmental degradation (Figure 5A–P).

**Figure 5.**
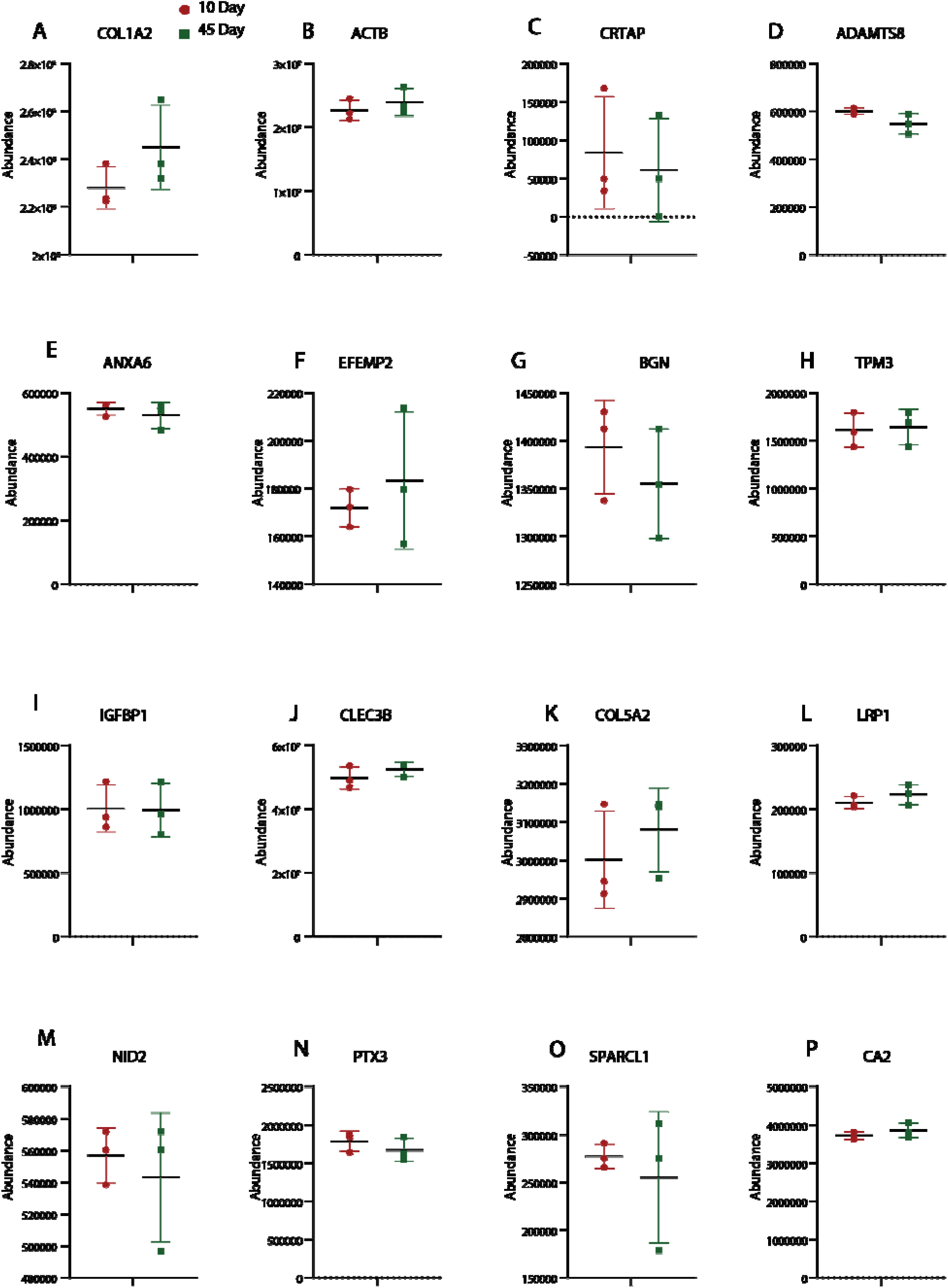
Targeted evaluation of selected temporally stable proteins with forensic relevance: Box plots showing quantitative abundance of selected stable proteins in Method 1 at 10 days and 45 days. Proteins were selected based on temporal stability and known relevance to bone extracellular matrix, remodeling, mineralization, or structural persistence. (A) COL1A2, (B) ACTB, (C) CRTAP, (D) ADAMTS8, (E) ANXA6, (F) EFEMP2, (G) BGN, (H) TPM3, (I) IGFBP1, (J) CLEC3B, (K) COL5A2, (L) LRP1, (M) NID2, (N) PTX3, (O) SPARCL1, and (P) CA2. Stable abundance across time points supports their candidacy as forensic persistence markers.

Several of the selected candidates were structural extracellular matrix proteins, including COL1A2, COL5A2, BGN, SPARCL1, NID2, EFEMP2, and CRTAP [30]. These proteins are involved in collagen fibril organization, matrix assembly, and tissue architecture, and their persistence is likely related to physical embedding within the mineralized bone matrix [31]. In particular, COL1A2, the major fibrillar collagen component of bone [30], remained highly abundant and stable over time, reinforcing its value as a benchmark skeletal persistence marker. Likewise, non-collagenous matrix proteins such as BGN and SPARCL1 may provide complementary markers with greater tissue specificity.

A second group of stable proteins was linked to matrix remodeling and extracellular regulation, including ADAMTS8, PTX3, CLEC3B, and LRP1 [32]. Although these proteins are functionally involved in tissue turnover, signaling, or proteolytic regulation, their maintained abundance suggests that selected regulatory proteins can remain detectable within the preserved bone microenvironment. Such markers may be informative not only for persistence, but also for assessing retained biological activity or matrix state. Additional candidates associated with bone physiology included CA2 and IGFBP1 [33]. Carbonic anhydrase II (CA2) is strongly associated with osteoclast-mediated bone resorption and mineral metabolism [34], while IGFBP1 is linked to insulin-like growth factor signaling pathways relevant to bone turnover. Their temporal stability suggests that physiological proteins involved in skeletal remodeling may also persist under early post-exposure conditions.

Interestingly, several intracellular structural proteins, including ACTB, TPM3, and ANXA6, also showed minimal temporal variation. Although less bone-specific than extracellular matrix proteins, these proteins may reflect durable cellular remnants protected within lacunae, membrane compartments, or protein complexes. Their reproducible detection may therefore support use as auxiliary normalization or preservation markers. To further investigate relationships among selected stable proteins, interaction network analysis was performed. This revealed dense connectivity centered around collagen and extracellular matrix-associated proteins (Supplementary Figure 4E), with major nodes including COL1A2, COL5A2, BGN, SPARCL1, and NID2. These findings suggest that temporal persistence is enriched among structurally integrated protein networks rather than random individual proteins.

Collectively, these data define a focused panel of temporally stable proteins with potential forensic value. Among these, COL1A2, COL5A2, BGN, SPARCL1, NID2, EFEMP2, and CA2 emerge as particularly promising candidates for future development as persistence biomarkers, normalization controls, or indicators of preserved skeletal material.

### 3.7: Temporally dynamic proteins define degradation-sensitive intracellular pathways

To identify proteins most susceptible to temporal change following environmental exposure, normalized abundance profiles were examined across all samples using unsupervised clustering. Heatmap analysis of the combined quantitative dataset identified multiple clusters with distinct abundance behaviors, including a prominent temporally responsive group designated Cluster 1 (Figure 6A). Cluster 1 was characterized by high abundance in 10-day samples from both Method 1 and Method 2, followed by marked reduction in 45-day samples. Re-clustering of this subset confirmed a highly coordinated temporal decrease across both extraction workflows (Figure 6B), indicating that the observed changes were reproducible and independent of extraction strategy. Violin plot analysis further demonstrated a clear downward shift in the abundance distribution of Cluster 1 proteins between 10-day and 45-day samples (Figure 6C). This pattern was consistent in both Method 1 and Method 2, supporting the conclusion that these proteins represent a temporally dynamic subset of the bone proteome that is sensitive to post-exposure degradation or progressive protein loss.

**Figure 6.**
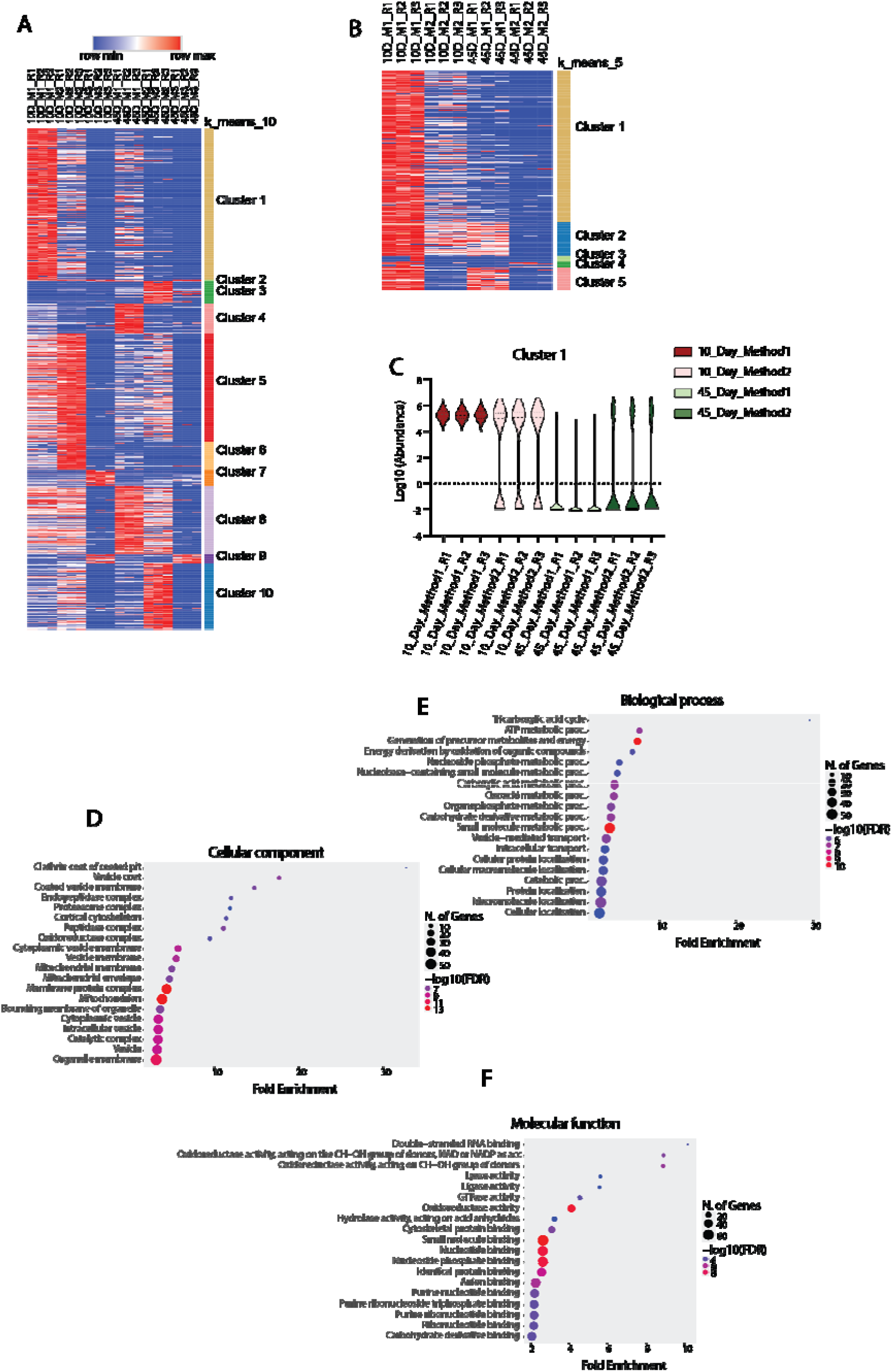
Identification and functional characterization of temporally dynamic proteins: **(A)** Heatmap of normalized abundance values across all quantified proteins showing ten major clusters with distinct temporal abundance profiles.**(B)** Expanded heatmap of Cluster 1 proteins demonstrating coordinated reduction in abundance between 10-day and 45-day samples across Method 1 and Method 2.**(C)** Violin plot showing log2-transformed abundance distribution of Cluster 1 proteins across methods and time points, illustrating temporal decline.**(D)** Gene Ontology cellular component enrichment analysis of dynamic proteins.**(E)** Gene Ontology biological process enrichment analysis of dynamic proteins.**(F)** Gene Ontology molecular function enrichment analysis of dynamic proteins.

#### 3.7.1: Dynamic proteins are enriched in vesicular, mitochondrial, and membrane-associated compartments

To determine the biological identity of temporally dynamic proteins, Gene Ontology enrichment analysis was performed. Cellular component analysis revealed significant enrichment for vesicle, vesicle coat, clathrin coat of coated pit, coated vesicle membrane, cytoplasmic vesicle membrane, mitochondrial membrane, mitochondrial envelope, membrane protein complex, and organelle membrane terms (Figure 6D). These findings indicate that proteins undergoing temporal loss are predominantly associated with membrane-bound intracellular structures and trafficking compartments rather than structurally protected extracellular matrix environments.

#### 3.7.2: Dynamic proteins are linked to metabolism, transport, and catalytic activity

Biological process enrichment showed strong association with ATP metabolic process, tricarboxylic acid cycle, generation of precursor metabolites and energy, oxidative metabolism, nucleoside phosphate metabolism, intracellular transport, and vesicle-mediated transport (Figure 6E). Additional terms related to cellular localization and catabolic processes were also enriched. Molecular function analysis demonstrated overrepresentation of nucleotide binding, nucleoside phosphate binding, anion binding, hydrolase activity, oxidoreductase activity, ligase activity, lyase activity, GTPase activity, and protein complex binding (Figure 6F), consistent with proteins involved in active cellular metabolism and regulatory processes.

#### 3.7.3: Temporal instability likely reflects reduced structural protection

In contrast to the stable protein module enriched in extracellular matrix proteins, the dynamic subset was dominated by intracellular metabolic and membrane-associated proteins. These proteins are less likely to be embedded within the mineralized collagen matrix and may therefore be more exposed to proteolytic cleavage, oxidative damage, hydrolysis, or environmental stress during post-exposure aging. The preferential decline of mitochondrial, vesicular, and catalytic proteins suggests that cellular remnants within bone degrade earlier than matrix-associated structural proteins.

### 3.8: Representative temporally dynamic proteins highlight selective loss of intracellular pathways

To further illustrate the protein classes most affected by temporal exposure, representative proteins from the dynamic module were examined individually in Method 1 samples at 10 and 45 days. These proteins were selected based on strong differential abundance, consistent trends across replicates, and biological relevance to mitochondrial metabolism, glycolysis, protein turnover, membrane trafficking, cytoskeletal organization, and stress responses. In most cases, these proteins were readily detected at 10 days but were markedly reduced or nearly absent at 45 days (Supplementary Figure 5A–A2), confirming progressive loss of specific intracellular protein populations.

Several of the strongest declining candidates were mitochondrial proteins, including ACO2, COX4I1, OGDH, PDHA1, ATP5PO, and ATP5MF, which function in the tricarboxylic acid cycle, oxidative phosphorylation, and ATP generation (Supplementary Figure 5C, K, O, Q, S, Z). Their pronounced decline suggests that mitochondrial remnants within bone are highly susceptible to post-exposure degradation. Similarly, multiple metabolic enzymes involved in glycolysis and energy metabolism, including PFKM, ALDOC, GPI, PGAM1, ACADM, and ACADL, were strongly reduced over time (Supplementary Figure 5A, B, F, I, M, Y), indicating preferential loss of proteins associated with active cellular metabolism. Proteins involved in protein synthesis and turnover were also temporally sensitive. Representative examples included RPL24 and EIF3A, together with proteasome-associated proteins PSMD1, PSMD6, and PSMD14 (Supplementary Figure 5G, T, U, V, A1), consistent with progressive breakdown of translational and proteostasis machinery. In addition, several membrane trafficking proteins, including RAB2A, AP2M1, COPB1, and FLOT2, showed marked reduction (Supplementary Figure 5J, N, P, X), supporting early degradation of vesicular and membrane-associated systems. Structural proteins such as ACTN4, ACTR3, EZR, and MYO1B also declined with time (Supplementary Figure 5D, H, L, W), indicating that intracellular scaffolding proteins are less resistant to temporal exposure than extracellular matrix proteins. Likewise, stress- and damage-associated proteins including DIABLO, HMOX1, and STAT1 were substantially depleted at 45 days (Supplementary Figure 5E, R, A2).

Overall, these findings suggest that proteins associated with exposed intracellular compartments and metabolically active cellular systems are more vulnerable to environmental degradation, whereas structurally embedded matrix proteins show greater temporal persistence.

## 4: Discussion

The present study provides a systematic evaluation of bone protein extraction workflows together with temporal proteomic profiling to assess both analytical performance and protein persistence under environmentally exposed conditions. Although bone is widely recognized as one of the most durable biological tissues and is frequently used in forensic, archaeological, and anthropological investigations, understanding of how extraction methodology influences proteome recovery and how specific protein classes are retained or degraded over time remains limited. By integrating data-independent acquisition (DIA) based mass spectrometry with comparative extraction strategies, this study demonstrates that methodological choice strongly influences total protein recovery, reproducibility, and extracellular matrix coverage, highlighting sample preparation as a critical determinant of downstream proteomic interpretation. A key biological finding was the separation of the bone proteome into two major functional groups: temporally stable matrix-associated proteins and dynamic intracellular proteins that were more susceptible to abundance loss over time. Stable proteins were predominantly enriched for collagens, extracellular matrix regulators, and structural components, whereas temporally dynamic proteins were associated with mitochondrial metabolism, vesicular transport, catalytic activity, and other cellular compartments. These observations support the concept that proteins embedded within the mineralized collagen matrix are preferentially protected from environmental degradation, while exposed cellular remnants are more rapidly lost. Collectively, these findings have important implications for forensic proteomics, where persistent matrix proteins may serve as biomarkers of skeletal preservation and dynamic proteins may provide complementary indicators of postmortem change, while also establishing a broader framework for biomarker discovery and temporal assessment of aged skeletal remains.

A major finding of the present study was the pronounced effect of extraction methodology on protein recovery, proteome depth, and analytical reproducibility. Previous bone proteomics studies have consistently shown that the dense mineralized matrix of skeletal tissue presents substantial challenges for efficient protein solubilization, and that no single extraction workflow uniformly recovers all protein classes [19, 20, 22, 35]. In agreement with these observations, the three workflows evaluated here produced markedly different proteomic outputs, highlighting the importance of extraction chemistry in shaping the detectable bone proteome. Method 1 generated the broadest overall proteome coverage together with the highest overlap across replicates, suggesting efficient recovery of both extracellular and intracellular protein fractions. This likely reflects the combined benefit of effective demineralization and strong detergent-mediated solubilization, approaches that have previously been reported to improve access to proteins entrapped within hydroxyapatite-rich matrices and structurally complex tissues [36, 37]. Such broad-spectrum extraction strategies are particularly advantageous for discovery-based proteomics where maximal protein depth is desired. Method 2 yielded a lower total number of identified proteins but showed strong recovery of collagens and extracellular matrix-associated proteins. Similar trends have been reported in studies using chaotropic agents such as guanidine hydrochloride, which are effective in disrupting protein–protein interactions and solubilizing heavily crosslinked matrix proteins [22, 38, 39]. Given that collagens and associated non-collagenous proteins represent the dominant organic fraction of bone, this selective enrichment may be advantageous for studies focused on skeletal biology, matrix remodeling, paleoproteomics, or forensic persistence markers. In contrast, Method 3 showed substantially reduced protein recovery and lower reproducibility under the present conditions. Acid-based extraction workflows have been used historically for collagen-focused analyses and peptide recovery from mineralized tissues; however, harsh extraction environments can also lead to limited recovery of non-collagenous proteins [22, 35], partial protein degradation, or reduced compatibility with broad shotgun proteomic workflows. The relatively low performance observed here therefore suggests that such approaches may be less suitable when comprehensive proteome coverage is the primary objective. Collectively, these findings reinforce the concept that extraction strategy is a critical analytical variable rather than a routine preparative step. Workflow selection should therefore be aligned with study aims: global proteome discovery may benefit from broad lysis-based methods, whereas extracellular matrix-focused applications may benefit from chaotropic enrichment strategies. More broadly, the results highlight the need for standardized and purpose-driven extraction protocols in forensic and skeletal proteomics to improve comparability across studies.

A central finding of the present study was the identification of a temporally stable subset of bone proteins that remained quantitatively conserved between 10-day and 45-day exposure conditions, indicating relative resistance to early environmental degradation. Previous studies in forensic science, paleoproteomics, and archaeological biomolecule research have consistently shown that structural proteins associated with the bone extracellular matrix are among the most persistent molecular components recovered from aged skeletal remains [22, 35]. In agreement with these observations, the stable protein module identified here was strongly enriched for collagens, matrix-associated glycoproteins, adhesion molecules, and other extracellular structural proteins, supporting the concept that biochemical stability is closely linked to tissue localization and structural embedding within the mineralized matrix. Among the most notable stable proteins were COL1A2, COL5A2, BGN, SPARCL1, NID2, EFEMP2, and CRTAP, all of which play recognized roles in collagen fibril organization, extracellular matrix assembly, or skeletal tissue integrity [40–42]. Type I collagen, the dominant organic constituent of bone, has long been regarded as one of the most durable proteins in vertebrate tissues due to its extensive crosslinking and intimate association with hydroxyapatite crystals [43–45]. Likewise, non-collagenous matrix proteins such as biglycan and SPARCL1 have been implicated in matrix organization, mineralization, and cell–matrix signaling [46–48], suggesting that persistence may extend beyond collagens alone to include broader extracellular networks. The interaction analysis performed in this study further supports this view, demonstrating that many stable proteins belonged to interconnected matrix-centered functional modules rather than representing isolated resistant species. From a forensic perspective, these stable proteins may have several practical applications. First, they may serve as robust biomarkers for confirming skeletal tissue origin in degraded or fragmentary remains. Second, because their abundance remains comparatively conserved over time, they may provide useful internal normalization targets for quantitative proteomic studies of aged bone. Third, persistent extracellular proteins could assist in assessing preservation quality prior to downstream molecular analyses such as DNA recovery or isotopic testing. More broadly, the results support the emerging concept that structurally protected matrix proteins represent the most reliable long-term molecular record within bone, and therefore should be prioritized in future biomarker development for forensic and archaeological investigations.

In contrast to the temporally stable extracellular matrix proteins, the present study identified a distinct subset of proteins that showed marked abundance reduction between 10-day and 45-day exposure conditions, indicating increased susceptibility to temporal degradation. Previous forensic and decomposition studies have suggested that intracellular proteins are generally less persistent than structural matrix proteins because they are not protected by mineral embedding and remain more accessible to autolysis, microbial activity, oxidation, and environmental stress [8, 14, 49]. The findings of the current study strongly support this model, as the dynamic protein cluster was predominantly enriched for mitochondrial, metabolic, vesicular, translational, and membrane-associated proteins. Among the most affected proteins were mitochondrial enzymes and respiratory components such as ACO2, SDHA, OGDH, PDHA1, ATP5PO, ATP5MF, and COX4I1, together with metabolic enzymes including PFKM, ALDOC, GPI, PGAM1, ACADM, and ACADL. These proteins are central to oxidative phosphorylation, the tricarboxylic acid cycle, glycolysis, and energy production, and their decline suggests rapid loss of metabolically active cellular remnants following environmental exposure. Similar reductions in mitochondrial and metabolic proteins have been reported in other postmortem proteomic studies, where cellular energy systems are among the earliest pathways disrupted after death [13, 50]. Proteins involved in vesicle trafficking, translation, and proteostasis were also strongly represented within the dynamic module, including RAB2A, AP2M1, VPS35, COPB1, EIF3A, EIF3B, PSMD1, PSMD6, and PSMB5. These observations are consistent with progressive collapse of intracellular transport systems and protein quality-control machinery during tissue aging. Likewise, the reduction of cytoskeletal proteins such as ACTN4, ACTR3, EZR, and MYO1B suggests that intracellular structural networks are also vulnerable to prolonged exposure, despite partial stability of some highly abundant cytoskeletal proteins in other contexts. From a biological perspective, these results support a two-compartment model of bone protein preservation in which extracellular matrix proteins remain relatively protected, whereas residual cellular proteins undergo preferential degradation over time. From a forensic standpoint, temporally dynamic proteins may therefore provide valuable complementary markers for estimating postmortem change or skeletal preservation state, particularly when interpreted alongside persistent structural proteins. Future studies examining longer exposure intervals and different environmental conditions may help determine whether specific dynamic proteins follow predictable decay trajectories suitable for postmortem interval modeling.

Beyond their biological significance, the proteomic patterns identified in this study have several practical implications for forensic science and the analysis of skeletal remains. One of the most immediate applications is the use of temporally stable extracellular matrix proteins as markers of bone tissue identity and preservation quality. Proteins such as collagens and associated matrix components that remain consistently detectable despite environmental exposure may provide reliable molecular confirmation of skeletal origin in highly degraded, fragmented, or commingled remains where conventional morphological assessment is limited. In addition, because these proteins showed comparatively stable abundance profiles, they may serve as internal reference markers for normalization in quantitative proteomic workflows applied to aged bone samples. A second important application lies in the use of temporally dynamic proteins as indicators of postmortem change. The reproducible decline observed in mitochondrial, metabolic, trafficking, and intracellular proteins suggests that selective protein loss may reflect progressive tissue degradation. If validated across larger cohorts, extended postmortem intervals, and diverse depositional environments, such proteins could contribute to multi-marker models for estimating preservation stage or postmortem interval. This approach would complement existing forensic tools based on entomology, taphonomy, imaging, or nucleic acid degradation, particularly in cases involving skeletonized remains where traditional soft-tissue indicators are unavailable.The present findings may also have relevance for triaging downstream molecular analyses. For example, proteomic assessment of stable versus degraded protein signatures could help determine whether a skeletal specimen is suitable for DNA recovery, isotope analysis, or other destructive testing, thereby improving sample prioritization in forensic casework. Similar approaches may also prove valuable in archaeological contexts, where rapid evaluation of biomolecular preservation is often required before extensive sampling.More broadly, this study supports the growing potential of mass spectrometry-based forensic proteomics as a complementary tool in human identification and skeletal biology. With continued methodological standardization, expanded reference datasets, and validation across environmental scenarios, bone proteome signatures may become increasingly useful for objective assessment of preservation status, temporal change, and evidential value in forensic investigations.

Despite the promising findings of this study, several limitations should be considered. First, the experimental design examined a relatively limited temporal window between 10-day and 45-day exposure conditions, which captures early post-exposure changes but does not reflect the longer-term degradation processes often encountered in forensic or archaeological settings. Future studies extending sampling over months or years will be important to determine whether the stable and dynamic protein patterns observed here persist over longer timescales. Second, environmental exposure was assessed under a specific set of conditions, whereas factors such as temperature, humidity, soil chemistry, microbial activity, and burial context are known to substantially influence skeletal decomposition and biomolecule preservation. Validation across diverse environmental scenarios will therefore be essential. A further limitation is the likely restricted sample size and biological diversity of the analyzed specimens, which may not fully capture inter-individual variation related to age, sex, health status, or skeletal site. In addition, although the two higher-performing extraction workflows were investigated in detail, other extraction chemistries and analytical platforms may produce different recovery patterns. From an analytical perspective, quantitative proteomics is also influenced by peptide detectability, missing values, and ionization efficiency, particularly for lower-abundance proteins. Finally, while this study identifies candidate markers of temporal stability and degradation, their forensic utility requires validation in larger independent cohorts and under real-world decomposition conditions. Accordingly, the present findings should be viewed as a foundation for future investigation rather than definitive forensic biomarkers.

In conclusion, this study demonstrates that both extraction methodology and temporal exposure are major determinants of the detectable bone proteome. Comparative workflow analysis showed that protein recovery is strongly influenced by extraction chemistry, with Method 1 providing the greatest overall proteome depth, Method 2 offering effective recovery of matrix-associated proteins, and Method 3 showing limited performance under the conditions tested. These findings emphasize the importance of selecting extraction strategies according to analytical objectives and support the need for standardized protocols in skeletal proteomics (Figure 7). Beyond methodological optimization, the study provides evidence for a biologically meaningful two-layer model of bone protein preservation. Structurally embedded extracellular matrix proteins, particularly collagens and associated matrix regulators, displayed strong temporal persistence and resistance to environmental degradation. In contrast, intracellular proteins associated with mitochondria, metabolism, vesicular transport, and cellular activity showed marked abundance decline over time, consistent with preferential degradation of exposed cellular compartments (Figure 7). This distinction helps explain why some proteins remain detectable in aged skeletal remains while others are rapidly lost.From a forensic perspective, these results highlight stable matrix proteins as promising candidates for skeletal tissue verification, preservation assessment, and internal normalization, whereas dynamic intracellular proteins may provide complementary markers of temporal change. Collectively, the present work establishes an analytical and biological framework for future bone proteomics studies and supports the growing application of mass spectrometry-based approaches in forensic science, archaeology, and biomolecular investigation of degraded skeletal remains.

**Figure 7.**
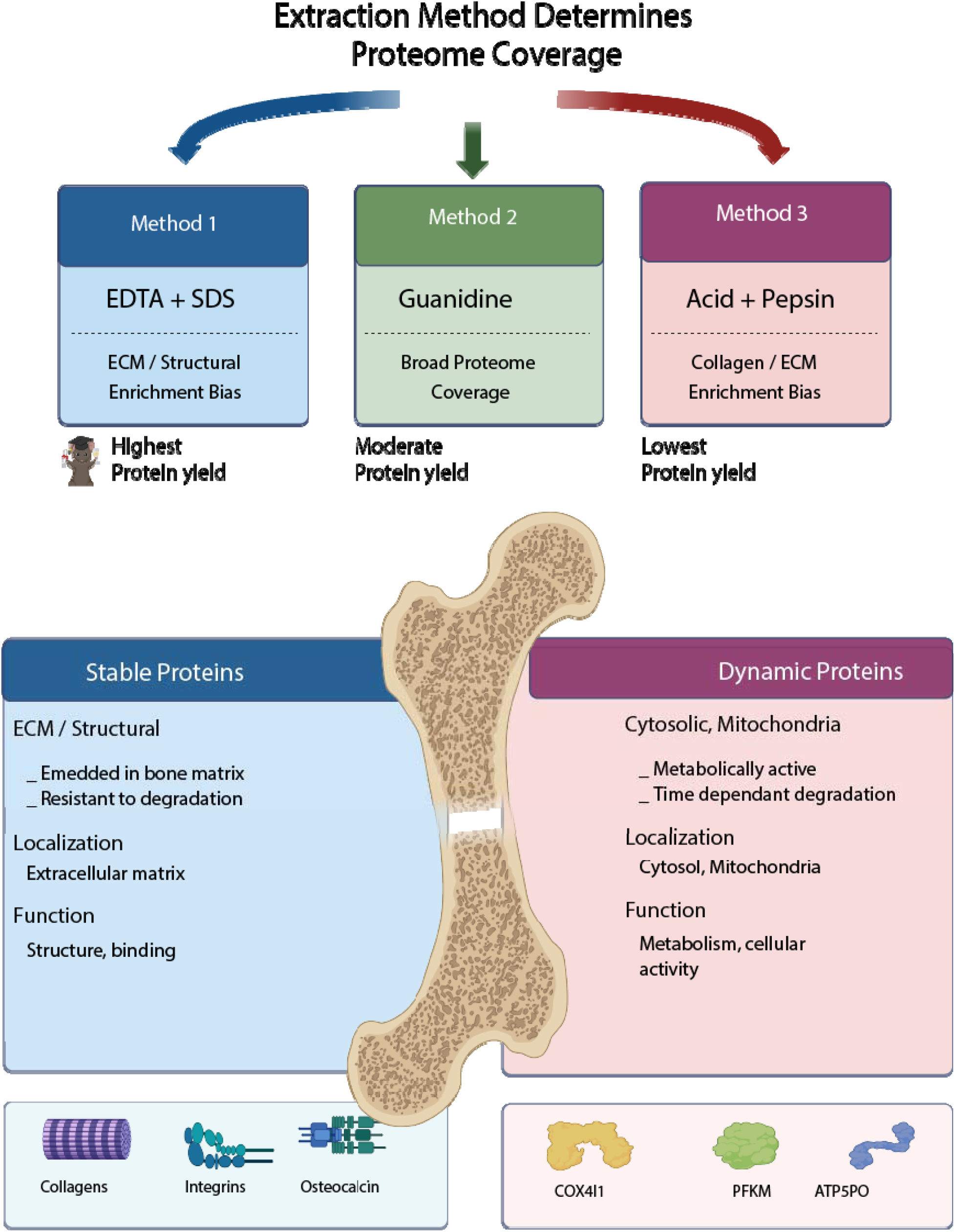
Summary model of extraction workflow performance and temporal stability of bone proteins: Schematic overview showing that extraction method influences detectable proteome coverage, under the tested conditions. Bone proteins were separated into two groups: stable proteins, mainly extracellular matrix/structural components resistant to degradation, and dynamic proteins, mainly cytosolic and mitochondrial proteins that declined over time.

## 5: Ethics approval and consent to participate

The Institutional Animal Ethics Committee approved the study with a reference number of YU/IAEC/P(L)10/2024.

## 6: Data Availability Statement

Data related to the study are provided in the manuscript and supplementary rest of the data and can be provided request.

## 9: Author Contributions

M.A.N conceived the idea, designed the experiments and interpretation of data, and critically reviewed and edited the manuscript. M.N.A conceived the idea, designed the experiments along with M.A.N. performed experiments and data analysis, drafted the manuscript, and prepared figures. N.C. Performed experiments, Proteomics sample preparation, acquired data, data analysis. S,A. Performed experiments, Proteomics sample preparation along with N.C. A.S Performed sample collection.

## 10: Acknowledgements

The authors gratefully acknowledge Yenepoya (Deemed to be University) for providing the infrastructure and state-of-the-art mass spectrometry facility required to conduct this study exclusively within our institution. We also acknowledge them for facility required and providing an opportunity to establish FACT. We acknowledge Dr. Vina Vaswani, Professor of Forensic Medicine and Director of the Center for Ethics. We acknowledge the Department of Forensic Medicine for its support. We also acknowledge the support provided by the Department of Biotechnology (DBT) through the National Facility grant under the project “Skill Development in Mass Spectrometry-based Metabolomics Technology BIC” (BT/PR40202/BTIS/137/53/2023).

## Abbreviations

DIA: Data-Independent Acquisition
LC-MS/MS: Liquid Chromatography-Tandem Mass Spectrometry
EDTA: Ethylenediaminetetraacetic Acid
SDS: Sodium Dodecyl Sulfate
GuHCl: Guanidine Hydrochloride
HCl: Hydrochloric Acid
TEAB: Triethylammonium Bicarbonate
DTT: Dithiothreitol
IAA: Iodoacetamide
FASP: Filter-Aided Sample Preparation
BCA: Bicinchoninic Acid (Assay)
UHPLC: Ultra-High-Performance Liquid Chromatography
MS: Mass Spectrometry
PCA: Principal Component Analysis
GO: Gene Ontology
FDR: False Discovery Rate
ECM: Extracellular Matrix

## Supporting information

All supplimentary data in doc file

