## Supplementary material for "Extraction-dependent bone proteomics reveals distinct stable and dynamic protein modules during early post-exposure degradation": All supplimentary data in doc file

Mohammad Nasir Ahmad, Ph.D.

Associate Professor

Department of Forensic Medicine,

Yenepoya medical college,

Yenepoya (Deemed to be University)

Mangalore 575018, India

**Authors**

Mohd. Altaf Najar

Nikita Choudhary

Sahla Abdulsalam

Aparna Sajeevan

Mohammad Nasir Ahmad

**Abstract**

Bone is a highly durable biological tissue widely used in forensic, archaeological, and anthropological investigations; however, efficient protein recovery and understanding of protein stability over time remain major challenges in skeletal proteomics. Here, we systematically evaluated three bone protein extraction workflows and integrated them with data-independent acquisition (DIA) mass spectrometry to assess proteome coverage, reproducibility, and temporal protein dynamics under environmentally exposed conditions. Comparative analysis demonstrated that extraction strategy is a primary determinant of detectable proteome composition. EDTA-based demineralization followed by SDS extraction provided the deepest proteome coverage and highest reproducibility, whereas guanidine hydrochloride extraction preferentially enriched collagen and extracellular matrix proteins. In contrast, acid-based extraction yielded limited protein recovery. Temporal profiling of bone samples collected at 10 and 45 days post-exposure revealed two distinct protein classes. A temporally stable module, enriched in collagens and extracellular matrix proteins including COL1A2, COL5A2, BGN, SPARCL1, and NID2, exhibited minimal abundance change, indicating resistance to environmental degradation. In contrast, temporally dynamic proteins, enriched in mitochondrial, metabolic, and intracellular pathways such as ACO2, OGDH, PDHA1, ATP5PO, and PFKM, showed marked decline over time. These findings support a two-compartment model of bone protein preservation in which matrix-embedded proteins are preferentially retained while exposed intracellular proteins undergo progressive degradation. Collectively, this study establishes an integrated framework linking extraction methodology with temporal proteome stability and identifies candidate markers for skeletal preservation assessment and temporal biomarker development in forensic and archaeological applications.

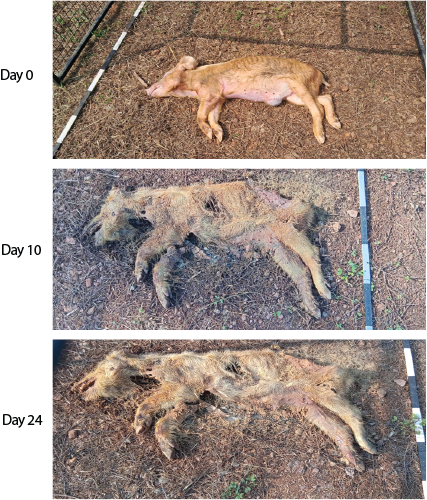

**Supplementary Figure 1. Representative images of specimen decomposition over time:** Images show progressive morphological changes under environmental exposure at Day 0 (intact), Day 10 (moderate decomposition), and Day 24 (advanced decomposition with extensive tissue loss and skeletal exposure).

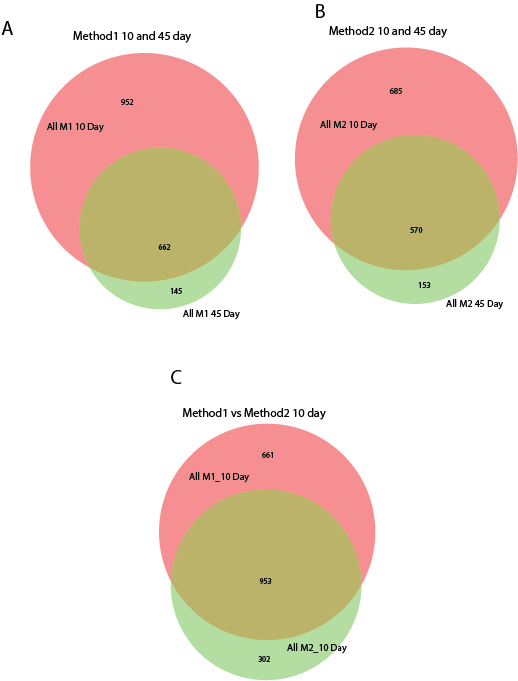

**Supplementary Figure 2. Protein overlap across methods and time points: (A)** Venn diagram showing overlap of identified proteins between **10-day and 45-day samples in Method 1**, highlighting shared and unique protein identifications across time points. **(B)** Venn diagram showing overlap of identified proteins between **10-day and 45-day samples in Method 2**. **(C)** Venn diagram comparing protein identifications between **Method 1 and Method 2 at 10-day**, illustrating shared core proteins and method-specific identifications. Numbers indicate unique and overlapping protein counts in each comparison. These analyses demonstrate a conserved core proteome across time points, along with method-dependent differences in protein recovery.

| Methods | Protein Accession | Protein | Gene | No of peptide |
| --- | --- | --- | --- | --- |
| M1 | A0A287AVY2 | procollagen galactosyltransferase | COLGALT1 | 1 |
|  | A0A480UHX7 | Procollagen C-endopeptidase enhancer 1 | PCOLCE | 17 |
|  | A0A481A6Z6 | Collagenase 3 | MMP13 | 22 |
|  | A0A287AFT5 | Collagen type XV alpha 1 chain | COL15A1 | 5 |
|  | A0A4X1TLE8 | Collagen type XI alpha 1 chain | COL11A1 | 38 |
|  | A0A287A0A6 | Collagen type VI alpha 6 chain | COL6A6 | 1 |
|  | A0A286ZWS8 | Collagen type II alpha 1 chain | COL2A1 | 12 |
|  | A0A4X1TP54 | Collagen triple helix repeat containing 1 | CTHRC1 | 7 |
|  | A0A286ZLV2 | Collagen alpha-3(VI) chain | COL6A3 | 86 |
|  | A0A480NML8 | Collagen alpha-2(XI) chain | COL11A2 | 16 |
|  | A0A480W6C8 | Collagen alpha-2(VI) chain | COL6A2 | 23 |
|  | A0A480N5F4 | Collagen alpha-2(V) chain | COL5A2 | 19 |
|  | A0A480F6B6 | Collagen alpha-1(XIV) chain | COL14A1 | 2 |
|  | A0A480VCL3 | Collagen alpha-1(XII) chain | COL12A1 | 162 |
|  | A0A480QX91 | Collagen alpha-1(VI) chain | COL6A1 | 20 |
|  | A0A480J5F2 | Collagen alpha-1(V) chain isoform 1 preproprotein | COL5A1 | 27 |
|  | A0A287A007 | Collagen alpha-1(IV) chain | COL4A1 | 4 |
|  | A0A1S7J210 | Collagen alpha-1(I) chain preproprotein | COL1A1 | 42 |
|  | A0A1S7J1Y9 | Alpha2 chain of type I collagen | COL1A2 | 36 |
|  | A0A480M2Y7 | 72 kDa type IV collagenase (Fragment) | MMP2 | 7 |
|  | A0A480J5F2 | Collagen alpha-1(V) chain | COL5A1 | 25 |
|  | A0A286ZQ85 | Collagen type III alpha 1 chain | COL3A1 | 2 |
|  | A0A287APU2 | Ficolin (collagen/fibrinogen domain containing) 1 | FCN1 | 9 |
|  | A0A4X1U2D4 | Collagen type IV alpha 2 chain | COL4A2 | 1 |
| M2 | A0A1S7J1Y9 | Alpha2 chain of type I collagen | COL1A2 | 46 |
|  | A0A286ZQ85 | Collagen type III alpha 1 chain | COL3A1 | 2 |
|  | A0A286ZWS8 | Collagen type II alpha 1 chain | COL2A1 | 19 |
|  | A0A1S7J210 | Collagen alpha-1(I) chain preproprotein | COL1A1 | 50 |
|  | A0A287AFT5 | Collagen type XV alpha 1 chain | COL15A1 | 4 |
|  | Q6VFT9 | Adiponectin, C1Q and collagen domain containing | ACDC | 1 |
|  | A0A287A007 | Collagen alpha-1(IV) chain | COL4A1 | 9 |
|  | A0A480M2Y7 | 72 kDa type IV collagenase (Fragment) | MMP2 | 6 |
|  | A0A480QX91 | Collagen alpha-1(VI) chain | COL6A1 | 11 |
|  | A0A480N5F4 | Collagen alpha-2(V) chain | COL5A2 | 23 |
|  | A0A480UHX7 | Procollagen C-endopeptidase enhancer 1 | PCOLCE | 13 |
|  | A0A480W6C8 | Collagen alpha-2(VI) chain | COL6A2 | 15 |
|  | A0A480J5F2 | Collagen alpha-1(V) chain isoform 1 preproprotein | COL5A1 | 37 |
|  | A0A481A6Z6 | Collagenase 3 | MMP13 | 24 |
|  | A0A4X1T6H1 | Procollagen C-endopeptidase enhancer 2 | PCOLCE2 | 2 |
|  | A0A4X1TLE8 | Collagen type XI alpha 1 chain | COL11A1 | 44 |
|  | A0A4X1TP54 | Collagen triple helix repeat containing 1 | CTHRC1 | 7 |
|  | A0A4X1U2D4 | Collagen type IV alpha 2 chain | COL4A2 | 4 |
|  | A0A480NML8 | Collagen alpha-2(XI) chain | COL11A2 | 25 |
|  | A0A480VCL3 | Collagen alpha-1(XII) chain | COL12A1 | 158 |
|  | A0A8D0QJ44 | Collagen alpha-1(XXIV) chain | COL24A1 | 2 |
|  | A0A286ZLV2 | Collagen alpha-3(VI) chain |  | 60 |
|  | A0A480NML8 | Collagen type XI alpha 2 (Fragment) | COL11A2 | 2 |
|  | A0A286ZIL9 | Collagen alpha-1(XVIII) chain | COL18A1 | 2 |
|  | A0A4X1U499 | Collagen IV NC1 domain-containing protein | COL4A3 | 1 |
|  | A0A480F6B6 | Collagen alpha-1(XIV) chain | COL14A1 | 3 |
|  | A0A480J5F2 | Collagen alpha-1(V) chain | COL5A1 | 38 |
|  | A0A286ZWS8 | Fibrillar collagen NC1 domain-containing protein |  | 10 |
|  | A0A287A1X1 | Collagen type XIII alpha 1 chain | COL13A1 | 1 |
|  | A0A4X1SP94 | Collagen type XXII alpha 1 chain | COL22A1 | 1 |
| M3 | A0A1S7J1Y9 | Alpha2 chain of type I collagen | COL1A2 | 15 |
|  | A0A1S7J210 | Alpha1 chain of type I collagen | COL1A1 | 13 |
|  | A0A286ZWS8 | Collagen type II alpha 1 chain | COL2A1 | 1 |
|  | A0A480I0M5 | Collagen alpha-1(IV) chain | COL4A1 | 2 |
|  | A0A480J5F2 | Collagen alpha-1(V) chain | COL5A1 | 2 |
|  | A0A480N5F4 | Collagen alpha-2(V) chain preproprotein | COL5A2 | 3 |
|  | A0A480NML8 | Collagen alpha-2(XI) chain | COL11A2 | 1 |
|  | A0A4X1TLE8 | Collagen type XI alpha 1 chain | COL11A1 | 2 |
|  | A0A4X1UX90 | Collagen alpha-1(XII) chain | COL12A1 | 14 |
|  | A0A480UHX7 | Procollagen C-endopeptidase enhancer 1 | PCOLCE | 1 |

**Supplementary Table S1. Detailed list of collagen and collagen-associated proteins identified across methods**

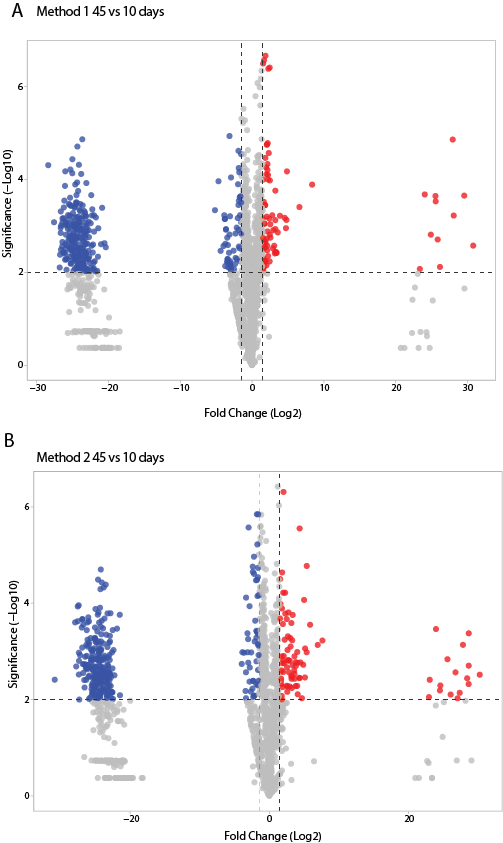

**Supplementary Figure3. Differential abundance analysis of temporal changes in the bone proteome:** Volcano plots showing differential protein abundance between 45-day and 10-day samples for Method 1 (A) and Method 2 (B). The x-axis represents log2 fold change (45-day vs 10-day), and the y-axis represents statistical significance (−log10 adjusted *p*-value). Proteins significantly increased at 45 days are shown in red, whereas proteins decreased at 45 days are shown in blue. Gray points indicate proteins not meeting significance thresholds.

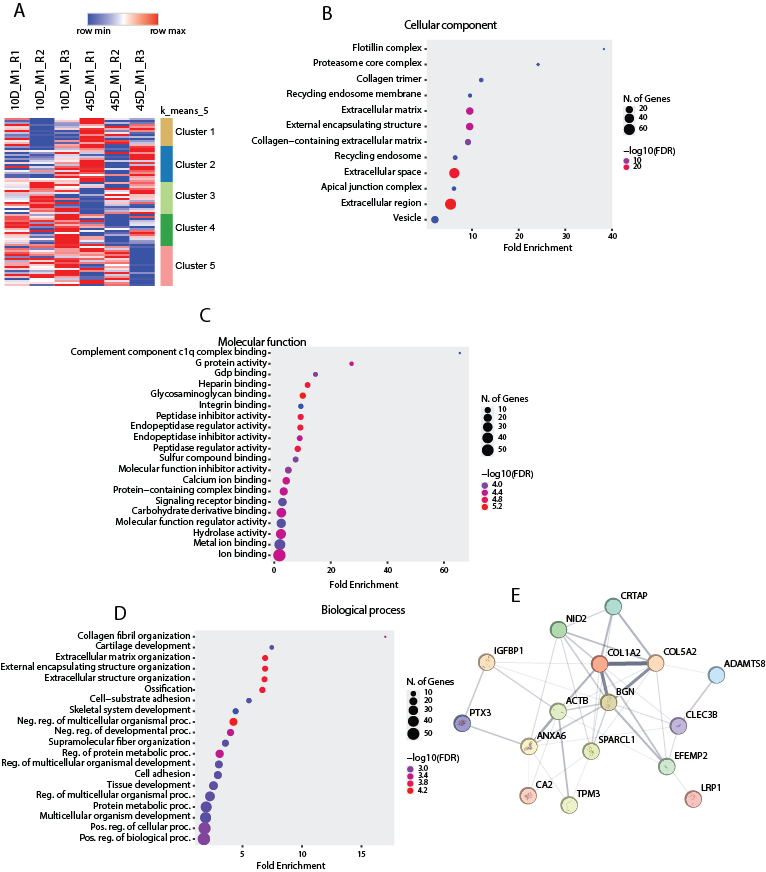

**Supplementary Figure 4. Identification of high-confidence temporally stable proteins:** (A) Heatmap showing abundance patterns of proteins meeting stringent temporal stability criteria across all samples: (B) Clustered visualization of the refined stable protein subset across extraction methods and time points. (C) Violin plot demonstrating conserved abundance distributions of high-confidence stable proteins. (D) Functional enrichment analysis of the refined stable protein panel highlighting extracellular matrix and structural protein categories **(E)** Protein–protein interaction network of selected high-confidence temporally stable proteins highlighting collagen-, extracellular matrix-, and structural protein-centered connectivity.

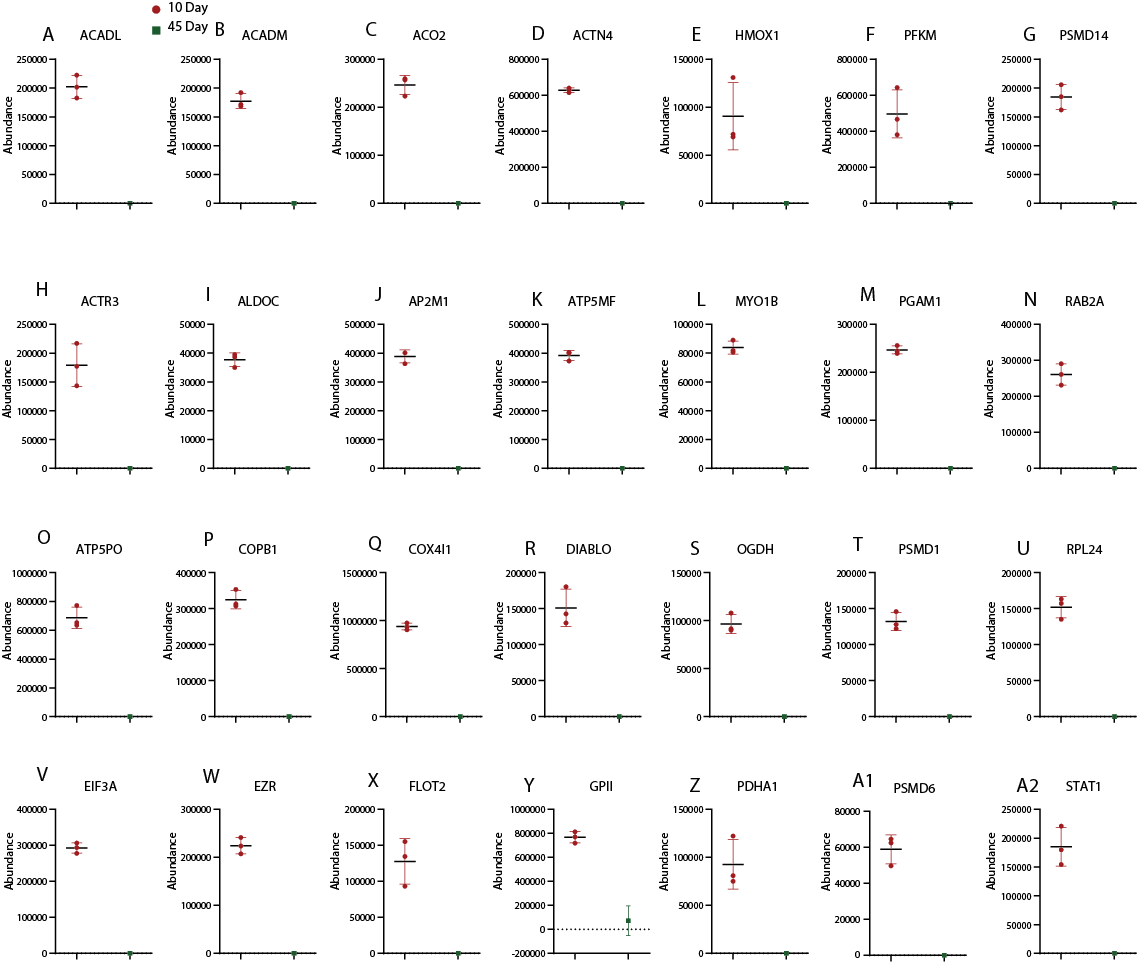

**Supplementary Figure 5. Representative temporally dynamic proteins identified in Method 1**: Abundance plots showing selected proteins with marked reduction between 10-day and 45-day samples in Method 1. Proteins were chosen from the dynamic cluster based on strong temporal decline and biological relevance to mitochondrial metabolism, glycolysis, protein turnover, membrane trafficking, cytoskeletal organization, and stress-response pathways. Consistent depletion at 45 days reflects increased susceptibility of exposed intracellular proteins to environmental degradation.
